# Genome-wide cell type-specific and sex-specific transcriptional dysregulation in the islet of Langerhans underlies islet dysfunction in Down syndrome-related diabetes

**DOI:** 10.64898/2026.03.12.711435

**Authors:** Cyrus R. Sethna, Maria del Carmen Mendoza Niemes, Bayley J. Waters, Matthew R. Wagner, Jacob M. Smith, Sutichot D. Nimkulrat, Julia Lemanski, Nicolas G. Pintozzi, Valentina Lo Sardo, Barak Blum

## Abstract

Individuals with Down syndrome (trisomy of human chromosome 21) are at a significantly higher risk of developing type 2 diabetes (T2D) than the general population. Systemic metabolic defects in Down syndrome have been linked to gene expression dysregulation in peripheral tissues like the liver, muscle, brain, and adipose. However, the contribution of gene expression dysregulation in the islets of Langerhans to the increased risk of T2D in Down syndrome has not been explored. Here we show that trisomic Ts65Dn mice, a common Down syndrome mouse model, are glucose intolerant and display reduced β-to-α cell ratio compared to disomic controls. Using single cell RNA sequencing on islets from Ts65Dn mice we found genome-wide, cell type-specific, and sex-specific transcriptional dysregulation in trisomic islets compared to controls. The Down syndrome-associated transcriptional signature revealed important islet defects, both at the cell autonomous level and at the whole-islet level, increasing T2D susceptibility. Our results put forth innate islet defects as a central underlying cause of Down syndrome-related T2D, warranting additional studies.

## Introduction

Down syndrome, caused by trisomy of human chromosome 21, is the most common chromosomal abnormality in the world, occurring approximately 1 in every 700 live births^1,2^. In addition to distinct physical features, intellectual disability and developmental delays, type 2 diabetes (T2D) has disproportionally higher incidence in individuals with Down syndrome^3^, who are four times more likely to develop T2D than the general population^4,5^. Moreover, individuals with Down syndrome are over 10 times more likely to develop T2D at younger ages (between 5-15 years of age), with Down syndrome-related T2D having less correlation with obesity than the general population^4^. The etiology of Down syndrome-related T2D is unknown and understudied^3^. Recent works have identified systemic metabolic defects in Down syndrome, which were attributed to gene expression dysregulation in the liver, muscle, hypothalamus, and adipose tissues^6–8^. However, the contribution of genetic dysregulation in the islets of Langerhans to Down syndrome-related T2D has not been established.

Several lines of evidence indicate that the increased T2D risk in individuals with Down syndrome also stems from inherent defects in the islets of Langerhans. First, individuals with Down syndrome display hypo-insulinemia and increased proinsulin-to-insulin ratio *in vivo* compared to age-matched controls, even when corrected for BMI and insulin resistance^9,10^. Second, *ex vivo* examination of islets from individuals with Down syndrome shows mitochondrial fragmentation, decreased insulin secretion, and an increase in islet amyloid polypeptide (IAPP) plaques^11^. To facilitate the investigation into how the presence of triplicated key regions in chromosome 21 affects overall cell and tissue physiology, several mouse models have been developed. In particular, Ts65Dn and Dp(16)1Yey mice, the two most common murine model of Down syndrome, carry an extra copy of a segment of mouse chromosome 16 that is homologous to a large section of human chromosome 21, and contains ∼57% and ∼65%, respectively, of the genetic material responsible for the Down syndrome phenotype^12,13^. Trisomic Ts65Dn and Dp(16)1Yey mice recapitulate Down syndrome phenotypes, including a predisposition to T2D^6,8,14^. Consistent with human-based data, Ts65Dn and Dp(16)1Yey mice show elevated fasting blood glucose levels and glucose intolerance despite having similar body weights, similar lean and fat mass, and (at least at a young age) similar insulin sensitivity to their disomic counterparts^8,14,15^. Additionally, trisomic Dp(16)1Yey mice were also reported to have reduced insulin content in their β cells, without reduction in β cell mass^14^.

Yet, with essentially no experimental studies examining the impact of trisomy 21 on pancreatic islets, innate islet abnormalities have not been previously implicated as a driving factor in Down syndrome-related T2D, and, as a consequence, therapeutic approaches to Down syndrome-related T2D targeting the islets have not been adequately explored. Here, we use the Ts65Dn mouse model to test the hypothesis that the predisposition to T2D in Down syndrome stems, at least in part, from inherent gene dysregulation in the islets of Langerhans.

## Results

### Disrupted Glucose Homeostasis Without Insulin Resistance in Young Trisomic

*Ts65Dn Mice* To explore islet function and glucose regulation in Down syndrome, we performed oral glucose tolerance tests (oGTT) and insulin tolerance tests (ITT) on young adult (6-8 weeks old) trisomic (ALT1) and disomic (WT) Ts65Dn mice. ALT1 male mice show significantly reduced glucose clearance following oGTT compared to WT littermate controls (Figure 1A; area under the cure (AUC), p=0.009, unpaired t-tests), consistent with previous reports^8,15^. Female ALT1 mice also show reduced glucose clearance after oGTT compared to WT controls, but this did not reach statistical significance (Figure 1B; AUC, p=0.0577, unpaired t-tests). Following ITT, male ALT1 mice showed identical glucose excursion curves relative to WT littermate controls, despite ALT1 mice having elevated glycemia set point (Figure 1C; area over the curve (AOC), p=0.7478, unpaired t-test). Female ALT1 mice also showed identical glucose excursion curves to those of littermate WT following ITT, but their set point glycemia was similar to the controls (Figure 1D). Importantly, ALT1 animals at all points had similar or slightly lower body weight than WT controls (Figure 1E-H). Thus, trisomic ALT1 mice have abrogated oGTT response compared to WT control littermates despite having normal insulin sensitivity and normal body weight, suggesting impaired islet function.

**Figure 1:**
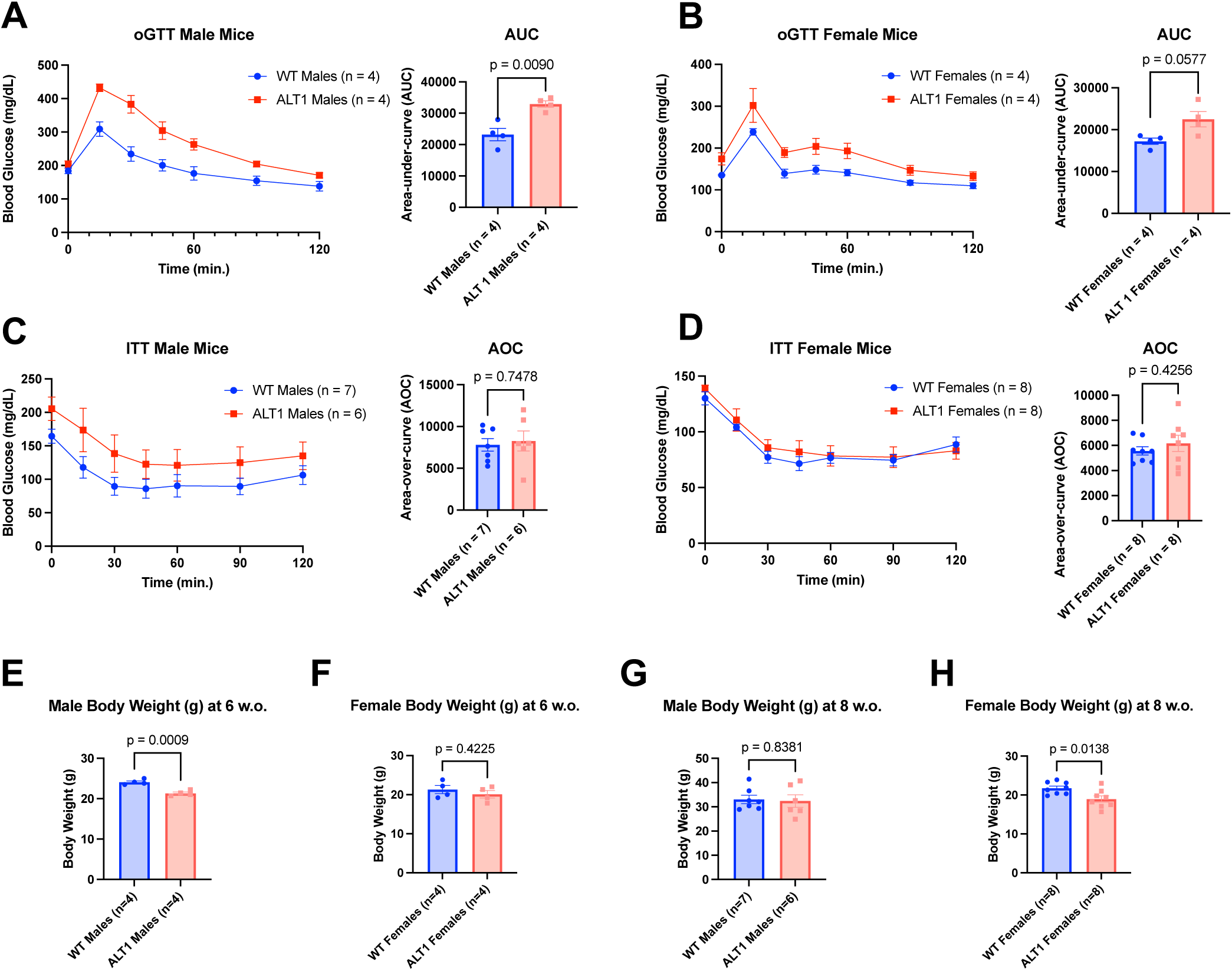
Disrupted glucose homeostasis without insulin resistance in young trisomic Ts65Dn mice. **(A)** Plasma glucose levels over 120 min after oral glucose tolerance test (oGTT) in 6-week-old male WT and ALT1 Ts65Dn mice (left) and corresponding area under the curve (AUC) (right; p=0.009, Welch’s t-test). **(B)** Plasma glucose levels over 120 min after oGTT in 6-week-old female WT and ALT1 Ts65Dn mice (left) and corresponding AUC (right; p=0.0577, Welch’s t-test). **(C)** Plasma glucose levels over 120 min after insulin tolerance test (ITT) in 8-week-old male WT and ALT1 Ts65Dn mice (left) and corresponding area over the curve (AOC) (right; Welch’s t-test, p=0.748). **(D)** Plasma glucose levels over 120 min after insulin tolerance test (ITT) in 8-week-old female WT and ALT1 Ts65Dn mice (left) and corresponding AOC (right; Welch’s t-test, p=0.4256). **(E-F)** Mouse body weights the time of the experiments in (A-D), respectively.

### Trisomic Ts65Dn Mice Have Altered Endocrine Cell-Type Composition

A few studies have shown that multiple tissues from individuals with Down syndrome, including the brain, blood, and heart, carry developmental defects resulting in altered cell type composition in the adult tissue^16–18^. No systematic and comprehensive analysis has been performed in the islet of Langerhans of individuals with Down syndrome. To assess islet endocrine cell types ratios in trisomic Ts65Dn mice compared to their disomic control littermates, we examined islet morphology and cellular composition in 9-week-old trisomic ALT1 mice and disomic WT littermate controls. The overall islet size and morphology were similar between trisomic ALT1 and WT controls for both males and females (Figure 2A,B). However, quantification of α, β, and δ cell proportions revealed slight alterations in endocrine cell type composition in both sexes. In males, ALT1 islets showed ∼5% more α cells and ∼5% fewer β cells, with no changes in δ cell proportion compared to WT controls (Figure 2C; ALT1 α: 20±8.9%, WT α: 14±5.2%, p=0.0002; ALT1 β: 58±13.9%, WT β: 65±8.3%, p=0.0031; ALT1 δ: 13±6.8%, WT δ: 13±6.5%, p=8.8573; unpaired t-test). In females, the effects were slightly larger: ALT1 islets contained ∼8% more α cells and ∼7% fewer β cells, with δ cell proportions remaining unchanged (Figure 2D; ALT1 α: 22±7%, WT α: 14±7.4%, p<0.0001; ALT1 β: 53±1%, WT β: 59±1.3%, p=0.0311; ALT1 δ: 17±8.3%, WT δ: 19±1.1%, p=0.3859; unpaired t-test). Despite these proportional shifts, analysis of the insulin-positive area revealed no differences in total β cell area between trisomic and disomic mice of either sex, suggesting possible compensatory hypertrophy of β cells (Figure 2E).

**Figure 2.**
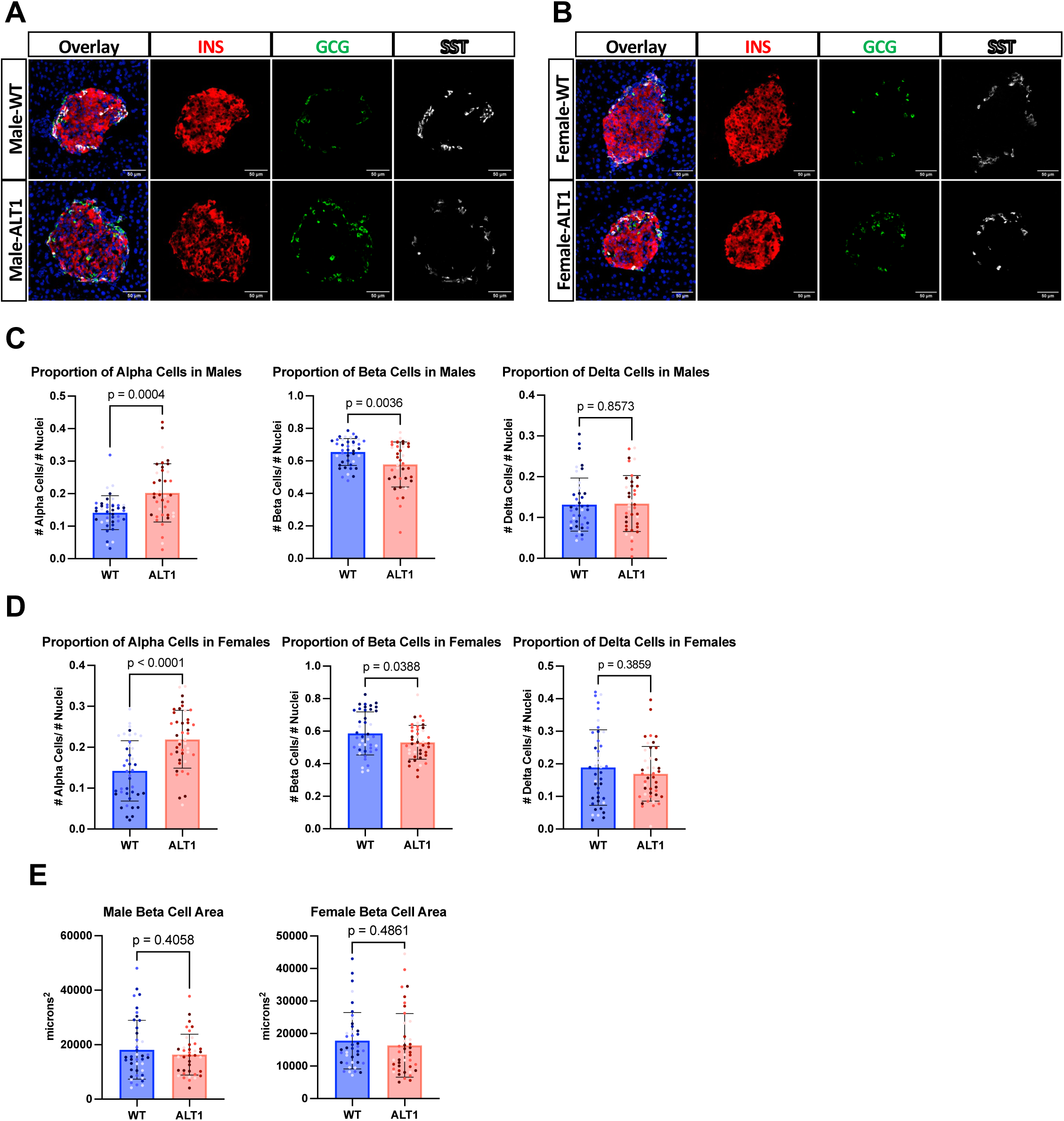
Trisomic Ts65Dn mice have altered endocrine cell-type composition. **(A-B)** Immunofluorescence staining for β cells (Insulin, red), α cells (Glucagon, green), and δ cells (Somatostatin, white) in islets of 9-week-old male **(A)** and female **(B)** WT and ALT1 Ts65Dn mice. Blue: DAPI. **(C-D)** Endocrine cell-type proportions in islets of 9-week-old male (C) and female (D) WT and ALT1 Ts65Dn mice (unpaired t-test; p-values indicated on graphs). **(E)** Average insulin-positive area in islets of male (left) and female (right) 9-week-old WT and ALT1 Ts65Dn mice; no significant differences were observed. Data are presented as mean ± SEM; n=4 mice per group, 10 islets imaged per mouse.

### Genome-Wide, Cell Type-Specific, and Sex-Specific transcriptional Dysregulation in Islets of Trisomic Ts65Dn Mice

To define the transcriptional landscape of islets in Down syndrome, we performed single cell RNA sequencing (scRNA seq) on isolated islets from young (6-weeks old) trisomic ALT1 mice and disomic WT littermate controls (Figure 3A). After quality control filtering, a total of 45,321 high-quality cells across four groups were further analyzed (ALT1 females: n=2; 11,255 cells; WT females: n=2; 6,723 cells; ALT1 males: n=2; 11,776 cells and WT males: n=3; 15,567 cells). Graph-based clustering resolved 16 transcriptionally distinct pancreatic populations (Figure 3B); marker-guided annotation then delineated endocrine and non-endocrine compartments. We identified all four endocrine cell types (α, β, δ, and PP), as well as two different clusters of polyhormonal cells (Poly 1 and Poly 2) (Figure 3C). Cluster presence and relative abundance did not show notable differences between ALT1 and WT islets or between sexes (Figure 3D,E).

**Figure 3.**
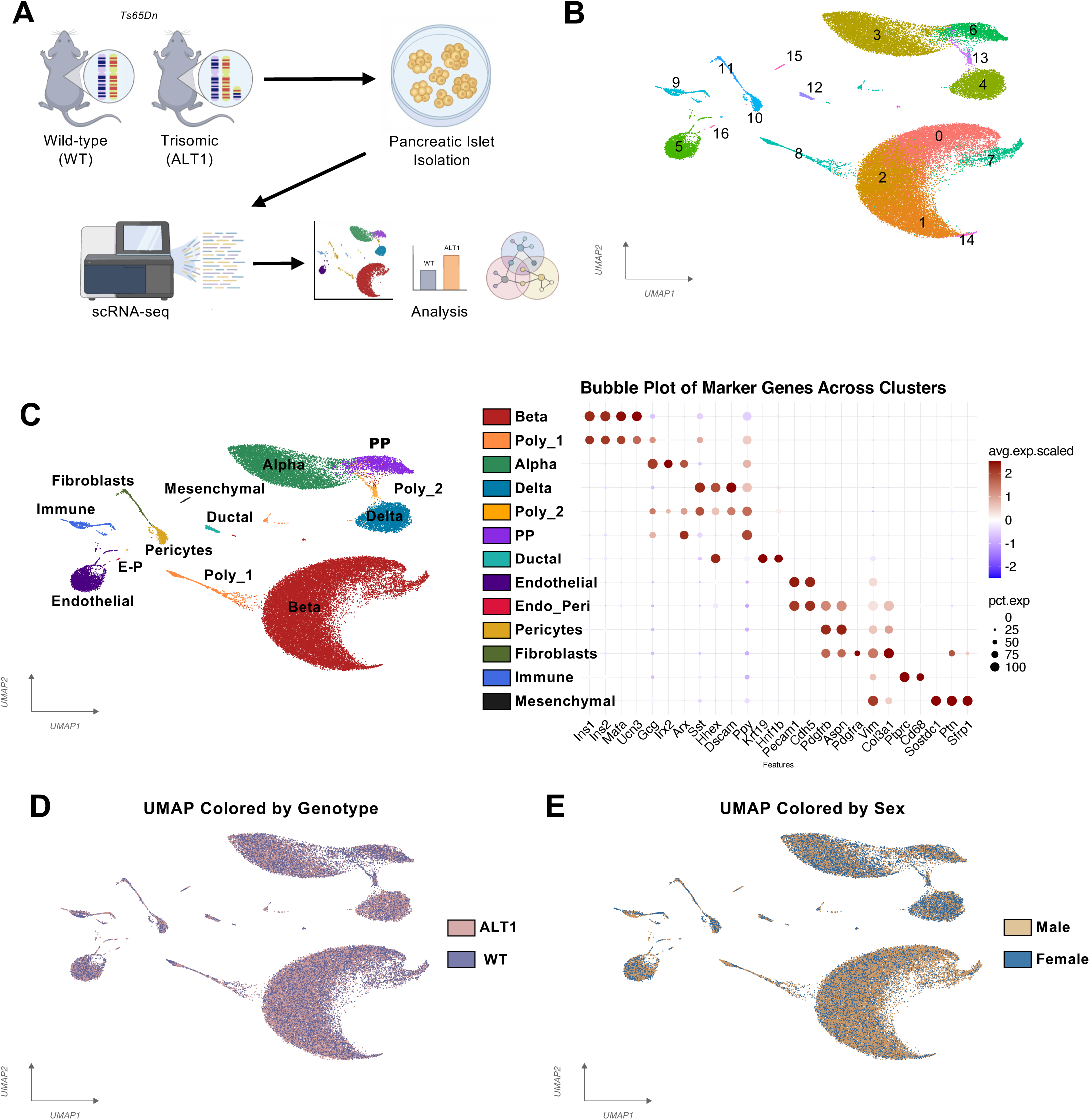
Single cell RNA-seq analysis of islets from disomic and trisomic Ts65Dn mice. **(A)** Schematic representation of the scRNA-seq workflow. **(B)** Graph-based clustering using Seurat. **(C)** Cell types assignment using canonical markers of pancreatic cell identity. **(D-E)** All identified cell types were visualized by genotype (**D**, WT in purple and ALT1 in pink) and sex (**E**, Male in yellow and Female in blue).

We conducted differential gene expression analysis between ALT1 and WT controls for each of the three major endocrine cell types. In male β cells we saw 1,339 upregulated genes, 103 of which were in the triplicated region, and 1,023 downregulated genes. Male α cells had 862 upregulated genes (76 triplicated), and 284 downregulated genes. Male δ cells had 358 upregulated (63 triplicated), and 184 downregulated genes. In females, we saw 1,438 upregulated (68 triplicated), and 2,172 downregulated genes in the β cells; 191 upregulated (42 triplicated), and 1,516 downregulated genes in the α cells; and 530 upregulated (53 triplicated) and 742 downregulated genes in the δ cells.

To assess whether DEGs were concentrated in specific chromosomal regions, we examined their distribution across the genome by chromosome. In both males and females, DEGs were evenly distributed across all chromosomes, with no evidence of enrichment on specific chromosomes. In agreement with previously reported data from other tissues^6,19–23^, we found that DEGs were spread across the entire genome, with the great majority of DEGs not located on the triplicated region (Figure 4A,B). This was consistent across endocrine cell types even at higher resolution, when plotted by gene start position within a chromosome (Supplementary Figure 1).

**Figure 4.**
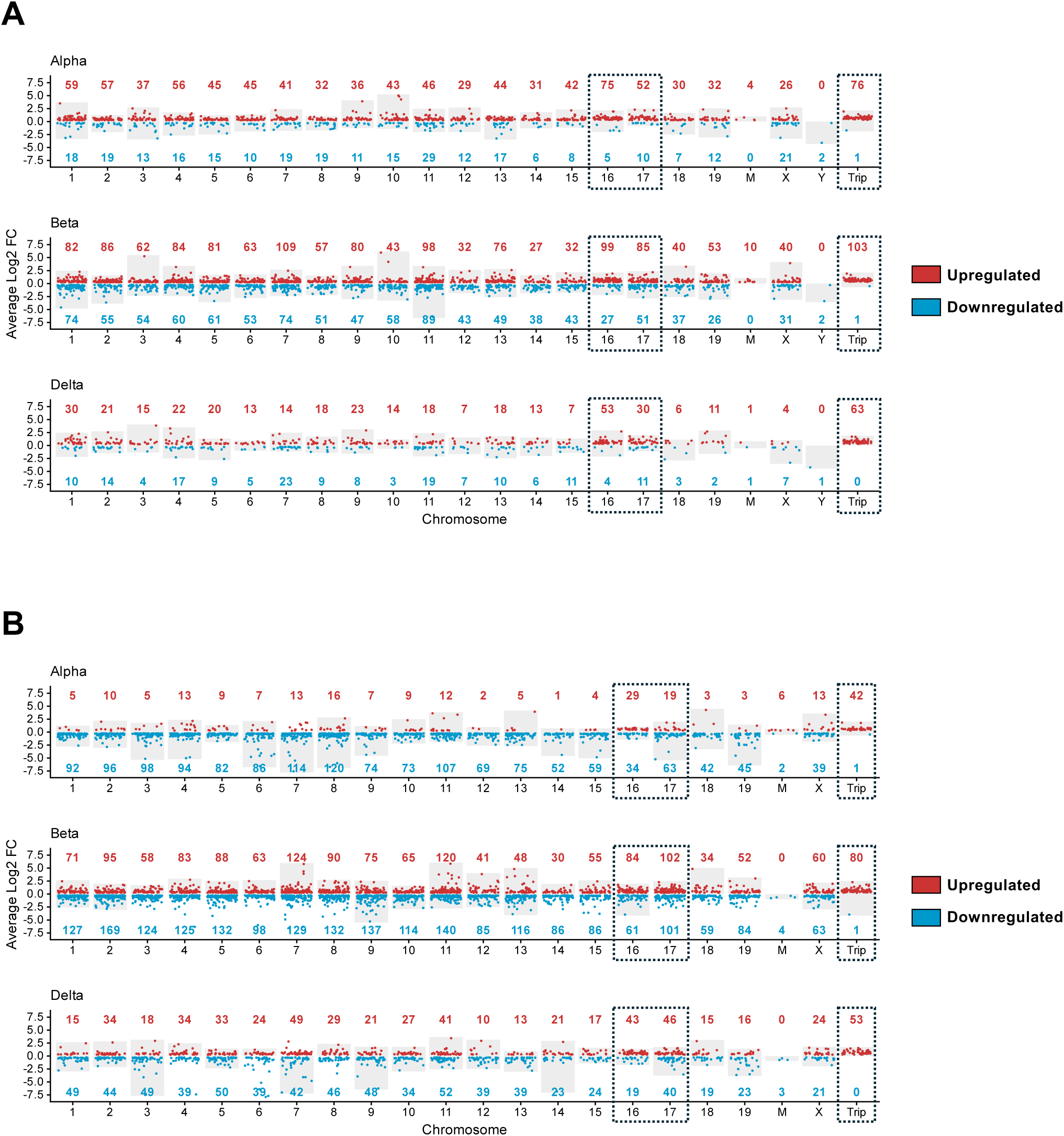
Genome-wide gene dysregulation in islets of trisomic Ts65Dn mice. **(A-B)** Chromosomal distribution of differentially expressed genes (DEGs) in endocrine cell types in males **(A)** and females **(B)** ALT1 mice compared to WT controls. Each point represents a gene, plotted by average log₂ fold-change (y-axis) and chromosome (x-axis). Upregulated genes in red and downregulated genes in blue. Numbers on red and blue above and below each chromosome represent the number of upregulated and downregulated DEGs in this chromosome, respectively. Dash-line box around chromosomes 16 and 17 indicated the presence of segmental triplication in these chromosomes. The dashed-lined box to the right (Trip) shows the relative expression of only the triplicated genes.

Remarkably, DEGs of males and females show little overlap in either upregulated or downregulated genes, except for the genes on the triplicated regions of chromosome 16, which were similarly upregulated in both sexes (Figure 5A). Moreover, very little overlap was observed among DEGs in α, β, or δ cells (Figure 5B). Together, these findings demonstrate that the gene dosage imbalance caused by Down syndrome drives a genome-wide transcriptional dysregulation in the islet that is sex- and cell type-specific.

**Figure 5.**
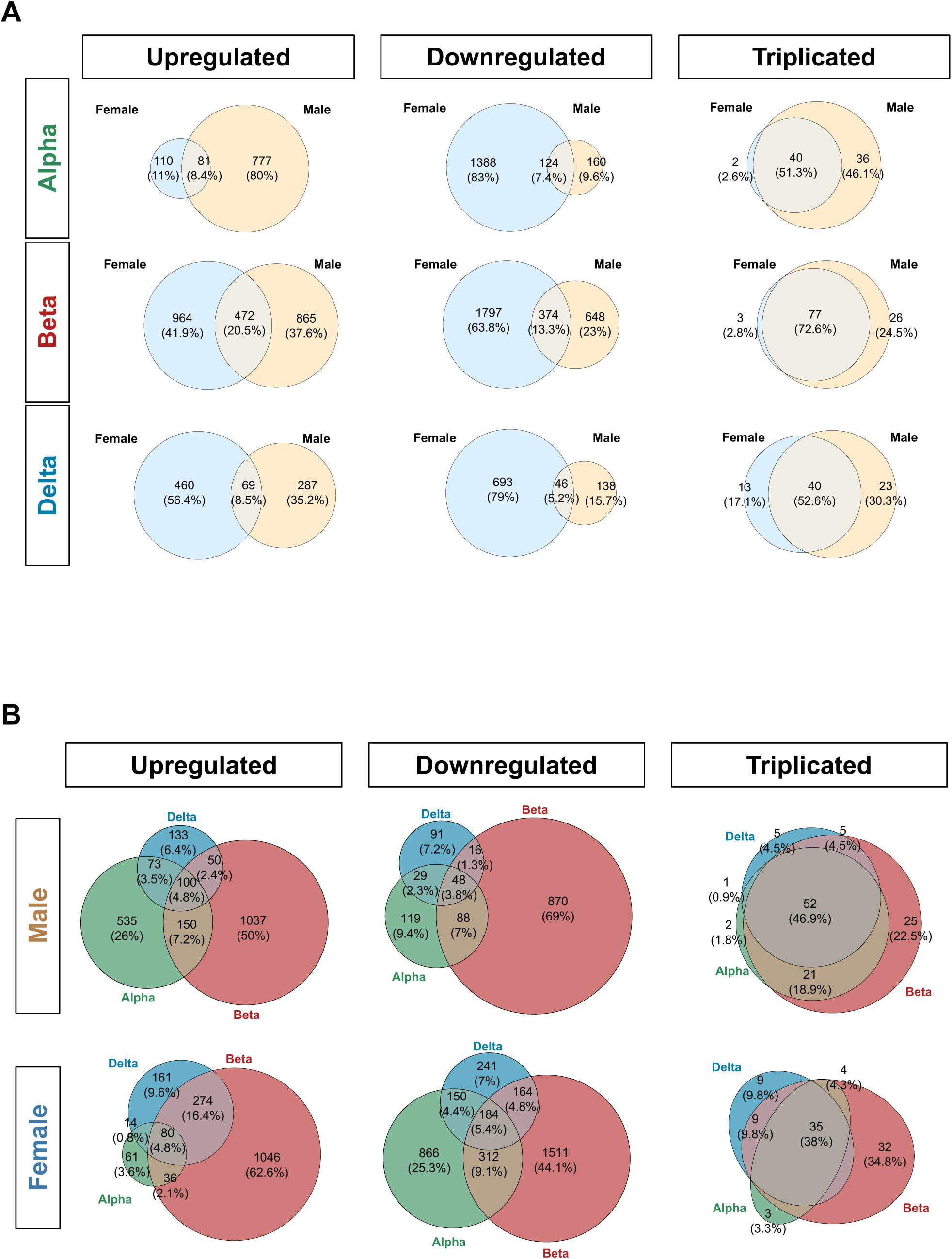
Cell type-specific and sex-specific gene dysregulation in islets of trisomic Ts65Dn mice. **(A)** Venn diagrams showing the overlap of differentially expressed genes (DEGs) between females (light blue, left) and males (yellow, right) within each cell type, for upregulated (left column; log₂ fold change [L2FC] > 0.25) and downregulated (center column; L2FC < −0.25) genes. Triplicated genes meeting the upregulation threshold are shown separately (right column). Gene counts and their percentages of the total are indicated within each set. **(B)** Venn diagrams showing the overlap of DEGs across cell types within each sex, using the same L2FC thresholds as in (A). β cells are shown in red, α cells in green, and δ cells in blue. All comparisons used an adjusted p-value threshold of <0.05.

### Overexpression of Triplicated Genes Related to T2D

We next examined the expression of genes triplicated in trisomic ALT1 mice. As expected, most (but not all) of the triplicated genes were upregulated ∼1.5-fold, consistent with gene dosage, in ALT1 compared to WT in three cell types (α, β, δ) in both males and females. The upregulated triplicated genes include several well-known diabetes-related genes whose function in the islets of Langerhans is documented, such as *Dyrk1a*, *Rcan1*, *Bace2*, *App*, *Bach1*, *Mpc1*, and *Sod1*. Notably, misexpression of any of these genes alone has been reported to negatively affect islet development, function, or compensatory response, suggesting that concurrent dysregulation of this set of genes in Down syndrome may be causal of increased and early-onset incidence of T2D (Refs.^15,24–33^) (Figure 6).

**Figure 6.**
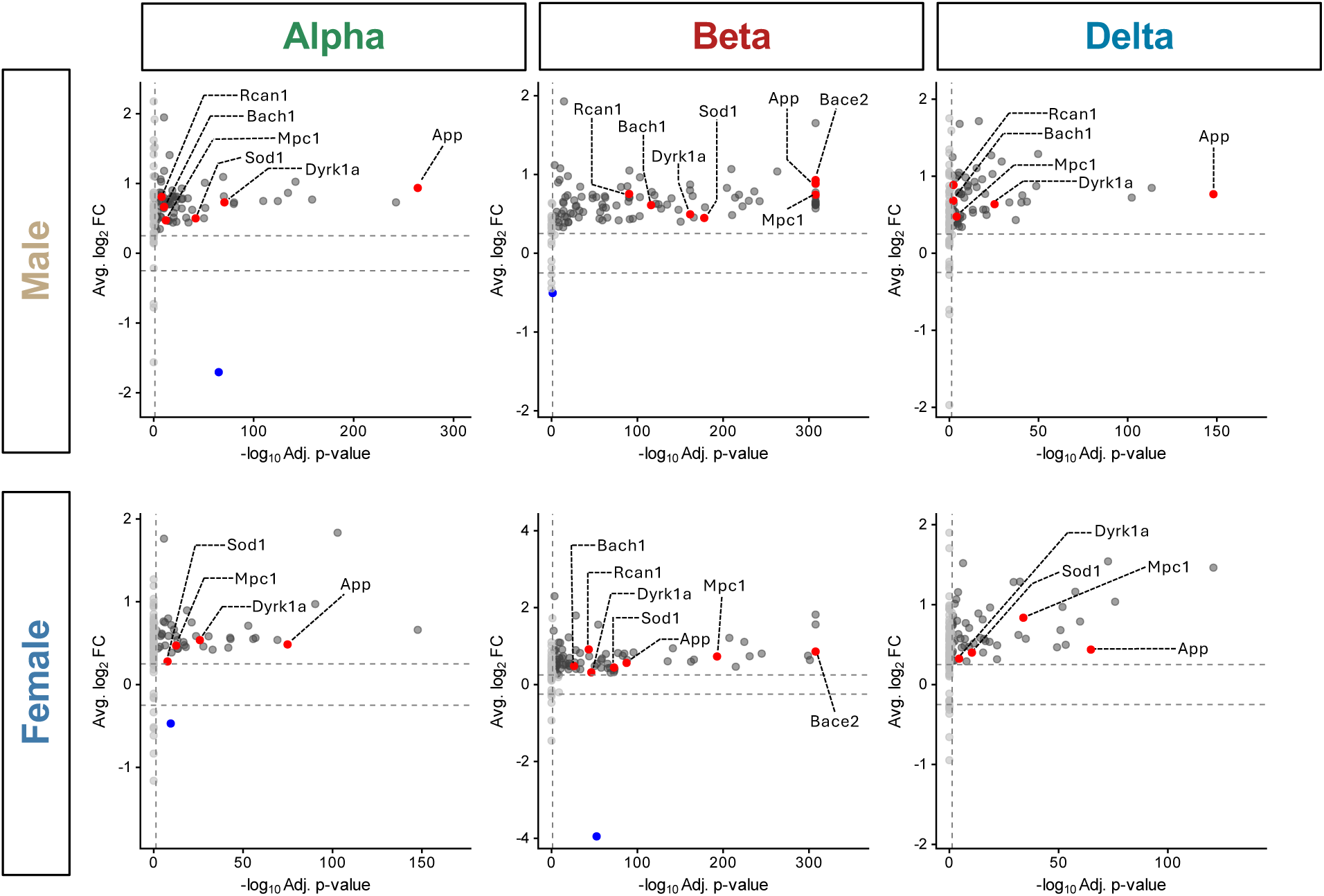
Overexpression of triplicated genes related to T2D. Volcano plots displaying differential expression of genes located on the triplicated genomic segment in Ts65Dn mice, stratified by sex and cell type. The top row represents male and the bottom row female mice. Columns correspond to α (left), β (middle), and δ (right) cells. Each point represents an individual gene, colored by significance and direction of expression: red indicates significantly upregulated genes discussed in the present study, dark grey indicates significantly upregulated genes not discussed, blue indicates significantly downregulated genes, and light grey indicates non-significant genes. Dashed grey line denotes the significance thresholds (L2FC = 0.25, p.adj<0.05).

As the overall transcriptional dysregulation in ALT1 islets spanned the entire genome and was vastly different between males and females and among the three major endocrine cell types, we performed Gene Ontology (GO) enrichment analysis on all upregulated and downregulated genes between ALT1 and WT islets across genotype, sex and cell types. After semantic simplification (see Methods), we analyzed the top 20 GO biological process (GO-BP) terms (comprehensive lists of all GO-BP terms and of all DEGs are shown in Supplementary Tables 1 and 2). We identify enriched GO-BP terms for each cell type and sex and highlight notable DEGs whose transcriptional dysregulation in the islets may explain the increased predisposition to T2D in individuals with Down syndrome.

### Transcriptional Dysregulation in Trisomic Ts65Dn Male β Cells

In male ALT1 β cells, the top 20 GO-BP terms of the upregulated genes largely fell into eight major categories: Endoplasmic reticulum (ER) stress and protein folding, Oxidative and chemical stress response, Hormone and nutrient response, Glycosylation and protein localization, Amino acid and protein metabolism, Vesicle transport secretion, Cytoskeleton adhesion, and Nucleotide ribonucleotide metabolism (Figure 7A). Notable upregulated genes related to ER stress and protein folding included ER chaperones and protein folding regulators such as *Dnajb9*, *Pdia6*, *Manf*, *Sdf2l1*, and *Erp44*, alongside canonical unfolded protein response (UPR) mediators *Atf6*, *Atf6b*, and *Creb3l2*. In addition, multiple components of the ER-associated degradation (ERAD) pathway were upregulated, including *Edem2*, *Ubxn4*, *Vcp*, *Optn*, and *Selenos*. Also included in this group was the upregulated triplicated gene *App*, an amyloid precursor protein whose upregulation has previously been implicated in islet dysfunction in diabetes^30^ (Figure 7B; grey). Genes related to oxidative stress and chemical stress response included the upregulation of *Rbp4*, an adipokine affecting β cell dysfunction in diabetes^34,35^; the triplicated genes *Rcan1* and *Sod1*; as well as the disallowed gene *Aldh1a3*, suggested to be a marker of β cell dedifferentiation^36,37^ (Figure 7B; dark blue). Within the hormone and nutrient response group, we saw an increase in vesicle priming, docking, fusion, and exocytosis genes such as *Stxbp5l*, *Unc13b*, *Rab3a*, *Rab13*, and *Doc2b*. The triplicated genes *Kcnj6*, encoding a G protein-activated inwardly rectifying potassium channel (GIRK2) subunit^38,39^, and *Synj1*, known to play a role in secretion granule recycling^40^, were also included in this group. Similarly, *Adora2a*, encoding a G protein-coupled receptor (GPCR) that modulate adenosine signaling in β cells^41^ was increased. Signaling modulators *Egfr* and *Pparg*, as well as granins *Chga* and *Vgf*, were also increased (Figure 7B; red). Notable upregulated genes related to glycosylation and protein localization included components of the oligosaccharyltransferase (OST) complex, *Rpn1*, *Rpn2*, *Stt3a*, *Ostc*, *Ddost*, and *Tmem258*. Genes involved in N-linked glycosylation were also increased including *Man2a1*, *Mgat2, Mgat3*, *Mgat5*, *Alg2*, *Alg8*, and *Alg14*. O-linked glycosylation enzymes were also upregulated, including *Galnt9*, *Galnt10*, *Galnt17*, and *Galnt18* (Figure 7B; orange). Finally, upregulated genes related to amino acid and protein metabolism included the triplicated genes *Dyrk1a* and *Bace2*, involved in regulating β cell proliferation and function^24–26,29^, as well as an increase in *Psen2*, which is another enzyme known for cleaving APP^42^. *Sirt4*, a mitochondrial enzyme known to repress the activity of glutamate dehydrogenase and thereby suppress insulin secretion in response to amino acids^43^, and *G6pdx*, the rate limiting enzyme of the pentose phosphate pathway^44,45^, were also increased. *Gad1*, which synthesizes GABA^46^, was also in this group (Figure 7B; light blue).

**Figure 7.**
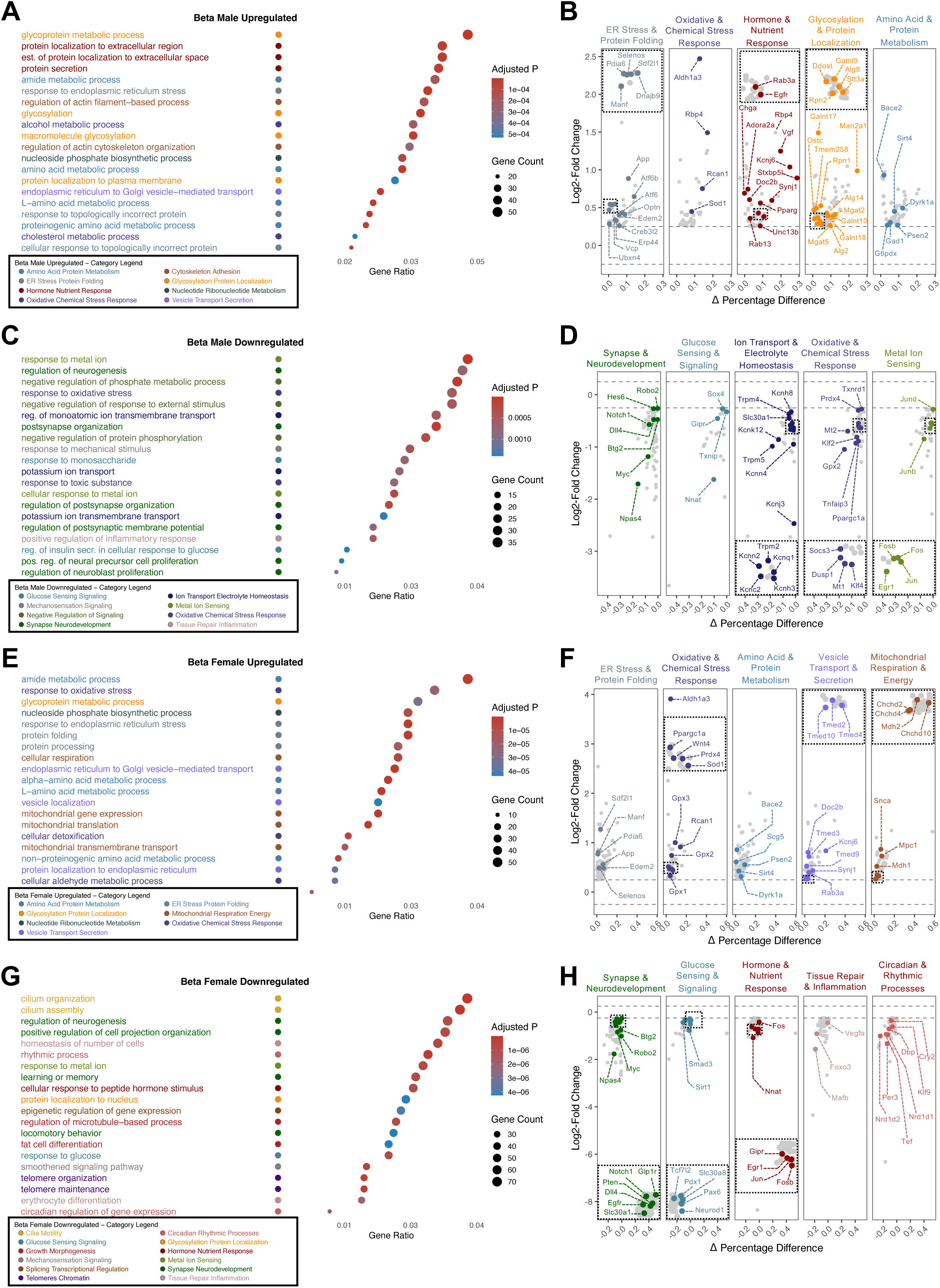
Transcriptional dysregulation in trisomic Ts65Dn β Cells. **(A-H)** Gene Ontology biological process (GO-BP) over-representation analysis (ORA) and differentially expressed genes (DEGs) in ALT1 β cells compared to wild-type controls, presented separately for males **(A-D)** and females **(E-H)**. **(A,E)** Top 20 over-represented GO-BP terms among upregulated genes, displayed as bar plots and colored by functional category. **(B,F)** Selected upregulated DEGs plotted as log₂ fold-change versus the difference in the fraction of expressing cells between ALT1 and WT (Δ fraction), with each sub-panel corresponding to a functional category and individual genes colored accordingly. **(C,G)** Top 20 over-represented GO-BP terms among downregulated genes, colored by functional category. **(D,H)** Selected downregulated DEGs plotted as in (B) and (F). Genes located within the Ts65Dn triplicated region and selected genes discussed in the text are labeled. Differential expression was determined using a threshold of |log₂ fold-change| ≥ 0.25 and adjusted p<0.05.

ALT1 male β cells compared to WT control showed a significant signature of downregulated genes, mostly involved in Synapse and neurodevelopment, Glucose sensing and signaling, Ion transport and electrolyte homeostasis, Oxidative and chemical stress response, Metal ion sensing, Mechanosensation signaling, Negative regulation of signaling, and Tissue repair inflammation (Figure 7C). Notable downregulated genes related to synapse and neurodevelopment included Notch signaling pathway components *Notch1*, *Dll4*, and *Hes6*, as well as of *Robo2*, a cell-surface receptor important for islet development and function^47–51^. *Btg2*, normally induced by incretin signaling through GLP-1 to promote insulin secretion^52^, was also reduced. *Npas4*, an important activity-dependent transcription factor in β cells^53–55^, was reduced. *Myc*, a driver of post-natal β cell proliferation also reduced, suggesting abnormal maturation-proliferation balance^56–59^ (Figure 7D; green). Notable downregulated genes related to glucose sensing and signaling included *Nnat*, which facilitates processing of nascent preproinsulin, and is important for β cell function^60–62^. Also in this group were *Sox4*, a key transcription factor regulating glucose tolerance through regulating β cell proliferation^63,64^, the thioredoxin-interacting protein *Txnip*, and, importantly, *Gipr*, which encodes the receptor for the incretin hormone GIP, and is involved in β cell function and survival^65,66^ (Figure 7D; red). Notable downregulated genes related to ion transport and electrolyte homeostasis included several members of the transient receptor potential melastatin (TRPM) family of non-selective cation channels (*Trpm2*, *Trpm4*, *Trpm5*) as well as numerous potassium channel genes (*Kcnc2*, *Kcnj3*, *Kcnn2*, *Kcnn4*, *Kcnq1*, *Kcnk12*, *Kcnh3*, *Kcnh8*), and a zinc transporter, *Slc30a1* (Figure 7D; dark navy blue). Notable downregulated genes associated with oxidative stress and chemical stress responses included the transcription factors *Klf2*, and *Klf4*, as well as antioxidant enzymes *Gpx2*, *Prdx4*, and *Txnrd1*. *Mt1* and *Mt2*, metallothioneins that regulate zinc availability and act as potent antioxidants^67^, were also included. Negative regulators of MAPK signaling (*Dusp1*) and NF-κB signaling (*Tnfaip3*), were also reduced, as was a negative regulator of the JAK-STAT pathway, *Socs3*. We also saw a downregulation of *Ppargc1a* (PGC-1α), an essential regulator of lipid metabolism^68^ (Figure 7D; dark blue). Finally, the downregulated metal ion sensing group was dominated by the immediate early genes (IEGs) *Egr1*, *Jun*, *Junb*, *Jund*, *Fos*, and *Fosb*, a group of rapid and transiently inducible genes that respond to metabolic signals in order to adapt to stress and maintain proper β cell function^69^ (Figure 7D; forest green).

### Transcriptional Dysregulation in Trisomic Ts65Dn Female β Cells

While female and male β cells showed some overlap in their dysregulated genes, most of the DEGs reflected sex-specific differences. Thus, the top 20 upregulated GO-BP terms fell into seven categories: ER stress and protein folding, Oxidative and chemical stress response, Amino acid and protein metabolism, Vesicle transport and secretion, Mitochondrial respiration, Glycosylation and protein localization, and Nucleotide ribonucleotide metabolism (Figure 7E). As in males, notable upregulated genes related to ER stress and protein folding included the triplicated gene *App*, the ERAD pathway components *Edem2* and *Selenos*, and the ER chaperone and protein folding regulators *Dnajb9*, *Pdia6*, *Manf*, and *Sdf2l1* (Figure 7F; grey). Similarly, notable upregulated genes associated with oxidative and chemical stress responses included the triplicated genes *Sod1* and *Rcan1*, and the disallowed gene *Aldh1a3*, which were also upregulated in male ALT1 β cells. However, *Prdx4*, *Ppargc1a*, and *Gpx2*, which were downregulated in the male β cells, are all upregulated in female β cells. Additionally, in female β cells, we saw an increase in another two glutathione peroxidase genes, *Gpx1* and *Gpx3*. Finally, *Wnt4*, which is heterogeneously expressed within β cell populations and marks more mature and functional β cells^70^, was increased (Figure 7F; dark blue). Related to amino acid and protein metabolism, there was upregulation of the triplicated genes *Dyrk1a* and *Bace2,* as well as the genes *Sirt4* and *Psen2*, as was also seen in the male β cells. In female β cells, we additionally saw an increase in *Scg5* (encoding the chaperone protein 7B2), which serves a critical role in the processing and maturation of proinsulin through the prohormone convertase PC2^71^ (Figure 7F; light blue). Notable genes related to vesicle transport and secretion included the triplicated genes *Kcnj6* and *Synj1*, as well as vesicle priming and exocytosis factors *Rab3a* and *Doc2b*, as seen in male β cells. Additionally, female β cells upregulate *Tmed2*, *Tmed3*, *Tmed4*, *Tmed9*, *Tmed10,* which serve as cargo receptors for protein trafficking from the ER to Golgi^72^ (Figure 7F; light purple). Notable genes related to mitochondrial respiration included the triplicated gene *Mpc1*, which encodes the mitochondrial pyruvate carrier 1, and facilitates pyruvate import into mitochondria for TCA cycle metabolism^73^, *Mdh1* and *Mdh2*, encoding the cytosolic and mitochondrial isoforms of malate dehydrogenase, respectively; *Snca* (encoding α-synuclein), which modulates potassium channels and negatively regulates insulin secretion^74^, *Chchd2* and *Chchd10*, which regulate cellular response to mitochondrial stress^75,76^, and *Chchd4*, the gene encoding the protein that imports them into the mitochondria^77^ (Figure 7F; dark red).

This transcriptomic signature was accompanied by strong downregulation of genes involved in Synapse and neurodevelopment, Glucose sensing and signaling, Hormone and nutrient response, Growth and morphogenesis, Tissue repair and inflammation, Circadian rhythmic processes, Glycosylation and protein localization, Cilia motility, Mechanosensing signaling, Metal ion sensing, Splicing and transcriptional regulation, and Telomeres and chromatin (Figure 7G). Notable downregulated genes related to synapse and neurodevelopment included Notch signaling components *Notch1* and *Dll4*, as well as *Btg2*, *Npas4*, *Myc*, *Robo2*, *Slc30a1*, and *Egfr*, all of which were also reduced in male ALT1 β cells. Interestingly, we also saw reduced expression of *Pten*, a negative regulator of PI3K/AKT signaling, whose deletion in adult β cells has been shown to induce regeneration^78^. *Glp1r*, the receptor for the incretin hormone GLP-1, which stimulates GSIS^66^, was also downregulated (Figure 7H; green). Notable genes related to glucose sensing and signaling included the transcription factors *Pax6, Neurod1*, and *Pdx1*, all of which are critical for maintaining β cell identity and function^79^, and *Smad3*, a downstream effector of the TGF-β signaling pathway, which is known to suppress β cell proliferation^80^. Importantly, *Slc30a8*, which transports zinc into insulin secretory vesicles, was also downregulated^81^. *Sirt1*, which regulates glucose metabolism and mitochondrial biogenesis^82–84^, and *Tcf7l2*, the most significant GWAS locus associate with T2D^85,86^, were also in this group of downregulated genes (Figure 7H; teal). Notable downregulated genes related to hormone and nutrient response included IEGs *Egr1*, *Jun*, *Junb*, *Fos*, and *Fosb*, alongside *Nnat*, and *Gipr*, all of which were also reduced in males ALT1 β cells (Figure 7H; red). Within the tissue repair and inflammation group we saw downregulation of transcription factors *Mafb* and *Foxo3*, as well as of *Vegfa*, normally secreted by β cells to promote islet vascularization^87^ (Figure 7H; pink). Finally, we saw downregulation of genes related to circadian and rhythmic processes, including of the circadian rhythm signaling regulators *Nr1d1*, *Nr1d2*, *Per3*, and *Cry2*; transcriptional regulator *Klf9*; and clock-output genes *Dbp* and *Tef* (Figure 7H; light red).

### Transcriptional Dysregulation in Trisomic Ts65Dn Male α Cells

Male ALT1 α cells exhibited a transcriptional profile that was distinct, and in some cases opposite from that of ALT1 β cells. The top 20 upregulated GO-BP terms in male ALT1 α cells fell into five categories: Synapse and neurodevelopment, Glucose sensing and signaling, Vesicle transport and secretion, Hormone and nutrient response, and Cytoskeletal adhesion (Figure 8A). Notable upregulated genes related to synapse and neurodevelopment included the triplicated genes *App, Rcan1, and Sod1* which are similarly upregulated in ALT1 β cells, but also *Robo2*, which is downregulated in male ALT1 β cells. *Cacna1a* and *Cacna1c*, which encode subunits of the P/Q type (CaV2.1) and L-type (CaV1.2) voltage-gated calcium channels, respectively, were also upregulated. The transcription factors *Foxa2* and *Klf7*, which regulate α cell and endocrine identity respectively^88,89^, were also included in this group (Figure 8B; green). Notable upregulated genes related to glucose sensing and signaling included several transcription factors that regulate α cell identity and function like *Arx*, *Rfx6*, and *Mafb*. Interestingly, we also saw an increased expression of *Neurod1*, which is suppressed in healthy mature α cells^90,91^, and of *Sox4*, known as a driver of endocrine lineage specification in pancreatic development^92,93^. In parallel, we also saw increases in *Pcsk2*, which encodes the enzyme Prohormone Convertase 2 (PC2) which processes proglucagon in mature glucagon^94^ (Figure 8B; teal). Notable genes associated with vesicle transport and secretion included the triplicated phosphatases *Synj1* and *Synj2*, which are vital for the endocytic cycle, impacting membrane recycling and trafficking^95,96^, *Prkca*, a protein kinase C isoform which translocases to the membrane to directly enhance glucagon release^97,98^, *Syt7*, a principal calcium sensor for glucagon secretion^99^, and *Unc13b*, a vesicle priming factor^100^ (Figure 8B; light purple). Related to the hormone and nutrient response category, notable upregulated genes included the triplicated potassium channel *Kcnj6*, alongside *Stxbp5l*, an inhibitor of insulin secretion in β cells^101^, and the pancreatic transcription factor *Hnf1b* (Figure 8B; red). Moreover, we found upregulation of genes related to cell adhesion like *Epha3*, *Sema3e*, *Ncam1*, *Cdh1*, and *Ntn4*, as well as the integrins *Itga6* and *Itgb5* (Figure 8B; orange).

**Figure 8.**
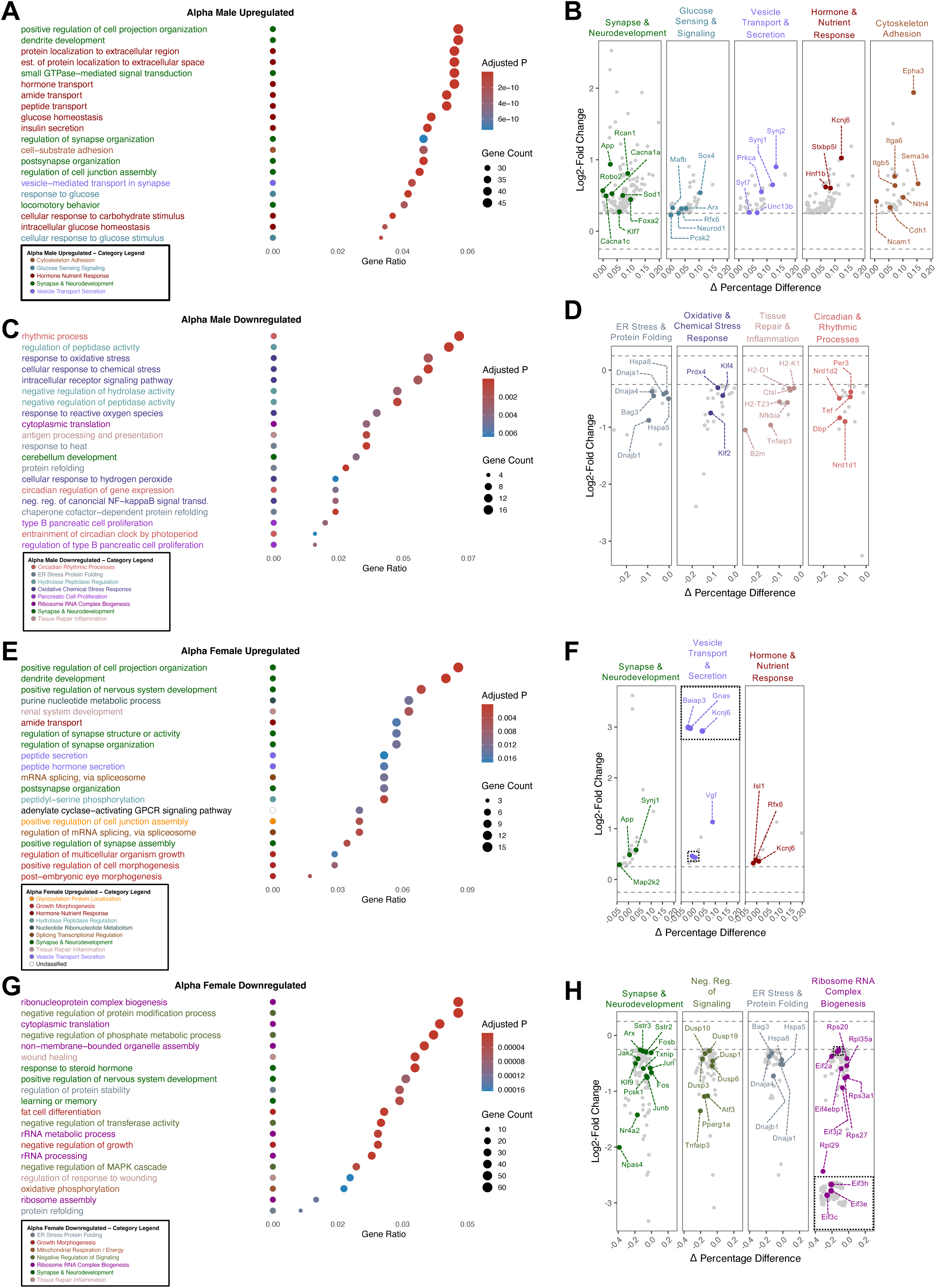
Transcriptional dysregulation in trisomic Ts65Dn α Cells. **(A-H)** GO-BP over-representation analysis and DEGs in ALT1 α cells compared to wild-type controls, presented separately for males **(A-D)** and females **(E-H)**. Layout and axes as in Figure 7.

The top downregulated 20 GO-BP terms of male ALT1 α cells fell into eight main categories: ER stress and protein folding, Oxidative and chemical stress response, Tissue repair and inflammation, Synapse neurodevelopment, Hydrolase peptidase regulation, Pancreatic cell proliferation, Ribosome RNA complex biogenesis, and Circadian rhythmic processes (Figure 8C). Notable downregulated genes related to ER stress and protein folding included UPR and autophagy genes *Hspa5* (GRP78/BiP), *Hspa8* (HSC70) and *Bag3*, and the Hsp40 co-chaperones *Dnaja1*, *Dnajb1*, and *Dnaja4* (Figure 8D; grey). Notable downregulated gene associated with oxidative and chemical stress responses included the transcription factors *Klf2* and *Klf4* (thought to regulate α cell identity^102^), antioxidant enzymes *Prdx4* and *Txnrd1*, the metallothionein *Mt1*, the negative regulator of MAPK signaling *Dusp1*, and the of mitochondrial regulator *Ppargc1a* (PGC-1α) (Figure 8D; dark blue). Notable genes related to tissue repair and inflammation included genes responsible for antigen presentation through MHC class I, *B2m*, as well as several MHC class I genes themselves, including *H2-K1*, *H2-D1*, and *H2-T23*, as well as *Ctsl* (encoding the lysosomal cysteine protease Cathepsin L1); negative regulators of NF-κB signaling, *Tnfaip3* and *Nfkbia* were also included in this group (Figure 8D; pink). Finally, notable downregulated genes related to circadian and rhythmic processes included *Nr1d1*, *Nr1d2*, *Per3*, *Dbp*, and *Tef* (Figure 8D; light red).

### Transcriptional Dysregulation in Trisomic Ts65Dn Female α Cells

The top 20 upregulated GO-BP terms in females ALT1 α cells fell into nine categories: Synapse and neurodevelopment, Vesicle transport and secretion, Hormone and nutrient response, Glycosylation and protein localization, Growth morphogenesis, Hydrolase peptidase regulation, Nucleotide ribonucleotide metabolism, Splicing transcriptional regulation, and Tissue repair inflammation (Figure 8E). Notable upregulated genes related to synapse and neurodevelopment included the triplicated genes *App* and *Synj1*, as well as *Map2k2*, a kinase within the MAPK/ERK pathway (Figure 8F; green). Notable genes related to vesicle transport and secretion included the triplicated gene *Kcnj6*, the Gαs protein subunit *Gnas*, which regulates glucagon expression and secretion^103^, and *Vgf*, a prohormone with important roles in systemic metabolic regulation^104,105^. *Baiap3*, involved in the maturation of dense-core vesicles in endocrine cells^106^, was also increased (Figure 8F; light purple). Notable upregulated genes related to hormone and nutrient response included the transcription factors *Isl1*, which regulated α cell lineage differentiation^107^, and *Rfx6*, also critical for α cell development and function^108^. Additionally, we also saw an increase in *Pde1c*, a calcium/calmodulin-dependent phosphodiesterase that regulates cAMP signaling^109^ (Figure 8F; red).

The top 20 downregulated GO-BP terms in female ALT1 α cells fell into seven categories: Synapse and neurodevelopment, Negative regulation of signaling, ER stress and protein folding, Ribosome and RNA complex biogenesis, Growth morphogenesis, Mitochondrial respiration, and Tissue repair inflammation (Figure 8G). Notable genes related to synapse and neurodevelopment included several IEGs (*Fos*, *Fosb*, *Jun*, *Junb*, and *Npas4*), as well as signaling regulators *Klf9*, *Jak2*, and the nuclear receptor *Nr4a2*, which regulates GLP-1 production in α cells^110^. We also saw downregulation of *Pcsk1* (PC1/3), which in α cells processes proglucagon into GLP-1, *Txnip*, whose deletion in α cells has been shown to improve diabetes-associated hyperglucagonemia^111^, the Somatostatin receptors *Sstr2* and *Sstr3*, and the key α cell transcription factor *Arx* (Figure 8H; green). Notable downregulated genes related to negative regulation of signaling included *Tnfaip3* and *Ppargc1a*. We also saw a decrease in genes associated with negative regulation of MAPK signaling including *Dusp1*, *Dusp3*, *Dusp6*, *Dusp10*, and *Dusp19*. Finally, *Atf3*, a stress-inducible transcription factor which has been shown to regulate glucagon transcription under low-glucose^112^, was also reduced (Figure 8H; olive green). Similar to male ALT1 α cells, notable downregulated genes related to ER stress and protein folding included *Hspa5* (GRP78/BiP), protein quality control and autophagy genes *Hspa8* and *Bag3*, and the Hsp40 co-chaperones *Dnaja1*, *Dnajb1*, and *Dnaja4* (Figure 8H; grey). Finally, notable genes related to ribosome and RNA complex biogenesis included numerous ribosomal subunits from both the small (RPS) and large (RPL) complexes, namely *Rps3a1*, *Rps20*, *Rps27*, *Rpl29*, and *Rpl35a*. Translation initiation factors *Eif2a*, *Eif3c*, *Eif3e*, *Eif3h*, *Eif3j2*, and *Eif4ebp1*, were similarly reduced (Figure 8H; purple).

### Transcriptional Dysregulation in Trisomic Ts65Dn Male δ Cells

The top 20 upregulated GO-BP terms in male ALT1 δ cells fell into four categories: Synapse and neurodevelopment, Vesicle transport and secretion, Growth and morphogenesis, and Cytoskeleton adhesion (Figure 9A). Notable upregulated genes related to synapse and neurodevelopment included, as seen in other cell types, an increase in the triplicated genes *App*, *Synj1*, and *Rcan1*. The triplicated gene *Dscam*, an axon-guidance protein regulating synaptogenesis and arborization in neurons^113^, was also upregulated. We further saw upregulation of a component of the SWI/SNF chromatin remodeling complex, *Arid1b*, as well the P/Q- and L-type calcium channels *Cacna1a* and *Cacna1c* (Figure 9B; green). Notable upregulated genes related to vesicle transport and secretion included the ghrelin receptor *Ghsr*, which potently enhances glucose stimulated somatostatin secretion^114,115^, *Nrxn1*, *Mapk10*, and *Adam22*, a membrane-anchored protein involved in cell-cell and cell-matrix interactions^116^ (Figure 9B; light purple). Notable genes associated with growth and morphogenesis included the upregulation of the triplicated gene *Dyrk1a*, as well *Ctnnb1* and *Cdh1* (Figure 9B; cherry red).

**Figure 9.**
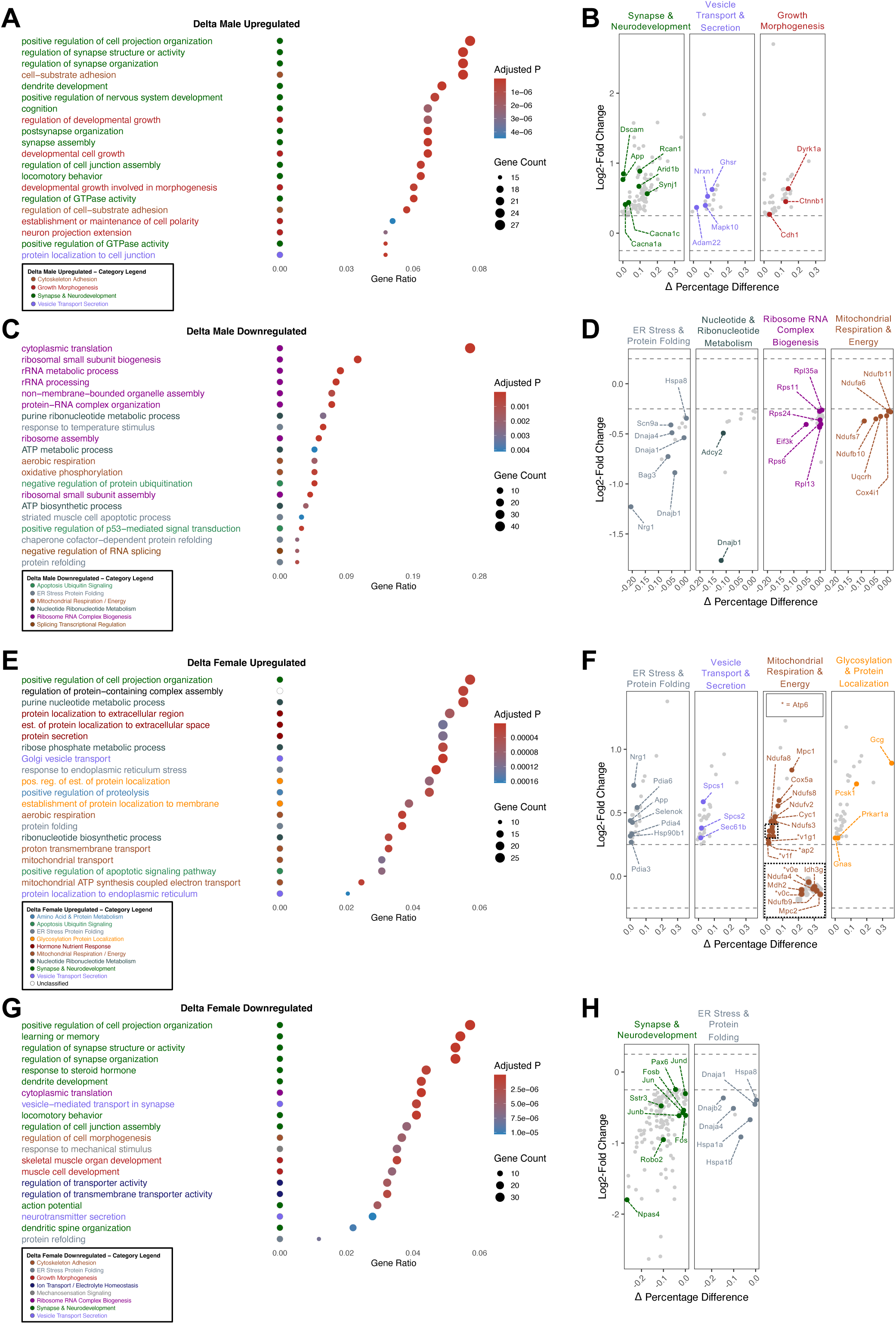
Transcriptional dysregulation in trisomic Ts65Dn δ Cells. **(A-H)** GO-BP over-representation analysis and DEGs in ALT1 δ cells compared to wild-type controls, presented separately for males **(A-D)** and females **(E-H)**. Layout and axes as in Figure 7.

Genes downregulated in male ALT1 δ cells fell into six main GO-BP categories: ER stress and protein folding, Nucleotide and ribonucleotide metabolism, Ribosome RNA complex biogenesis, Mitochondrial respiration and energy, Apoptosis ubiquitin signaling, and splicing transcriptional regulation (Figure 9C). Notable genes related to ER stress and protein folding included Hsp40 co-chaperones *Dnaja1*, *Dnajb1*, and *Dnaja4*, as well as genes involved in protein quality control and autophagy, *Hspa8* and *Bag3*. Further, *Scn9a,* a sodium channel, and *Nrg1*, a growth factor which signals thorough the ErbB receptors, were also downregulated (Figure 9D; grey). Notable genes related to nucleotide and ribonucleotide metabolism include *Pde8b*, a cAMP-specific phosphodiesterase that regulates the cAMP pool^117^, and *Adcy2*, encoding adenylate cyclase 2 which catalyzes the conversion of ATP into cAMP^118^ (Figure 9D dark grey). The downregulated Ribosome RNA complex biogenesis group encompassed multiple ribosomal subunits from the RPS and RPL complexes, including *Rps24*, *Rps6*, *Rps11*, *Rpl13*, and *Rpl35a*, as well as *Eif3k*, a component of the EIF3 complex (Figure 9D; purple). Finally, notable genes related to mitochondrial respiration and energy homeostasis included several subunits of the mitochondrial electron transport chain, including *Ndufb11*, *Ndufb10*, *Ndufs7*, and *Ndufa6* (complex I), *Uqcrh* (complex III), and *Cox4i1* (complex IV) (Figure 9D; dark red).

### Transcriptional Dysregulation in Trisomic Ts65Dn Female δ Cells

The upregulated genes in female ALT1 δ cells fell into the following main GO-BP categories: ER stress and protein folding, Vesicle transport and secretion, Mitochondrial respiration and energy, Glycosylation and protein localization, Amino acid protein metabolism, Apoptosis ubiquitin signaling, Hormone nutrient response, Nucleotide ribonucleotide metabolism, and Synapse neurodevelopment (Figure 9E). Notable genes related to ER stress and protein folding included upregulation of ER chaperones and folding regulators including *Manf*, *Hsp90b1*, *Pdia3*, *Pdia4*, and *Pdia6*, as well as ERAD pathway member *Selenok*. The triplicated gene *App* was also upregulated (Figure 9F; grey). Notable upregulated genes related to vesicle transport and secretion included *Sec61b*, as well as ER-resident signal peptidase subunits *Spc1* and *Spc2* (Figure 9F; grey light purple). Unlike in male ALT1 δ cells, we saw an upregulation of genes within the mitochondrial respiration and energy homeostasis group, including genes encoding subunits of the proton pump V-ATPase *Atp6ap2*, *Atp6v0c*, Atp6v0e, *Atp6v1f*, and *Atp6v1g1*. We also saw increased expression of genes encoding various mitochondrial respiratory chain complexes, including *Cox5a* (Complex IV) and *Cyc1* (Complex III), as well as members of the TCA cycle *Idh3g* and *Mdh2*. Notably, both subunits of the Mitochondrial Pyruvate Carrier *Mpc1*, a triplicated gene, and *Mpc2*, were upregulated. Further, unlike in the male ALT1 δ cells, Electron Transport Chain Complex I subunits *Ndufa4*, *Ndufa8*, *Ndufb9*, *Ndufs3*, *Ndufs8*, and *Ndufv2*, were also upregulated (Figure 9F; grey dark red). Within the upregulated glycosylation and protein localization group we saw an increase in *Pcsk1* and *Gcg*. Further, *Prkar1a*, a regulatory subunit of PKA, and *Gnas*, were also upregulated (Figure 9F; grey-orange).

The top 20 downregulated GO-BP terms in trisomic female δ cells fell into the following categories: Synapse and neurodevelopment, ER stress and protein folding, Cytoskeleton adhesion, Growth morphogenesis, Ion transport electrolyte homeostasis, Mechanosensation signaling, Ribosome RNA complex biogenesis, and Vesicle transport secretion (Figure 9G). Notable downregulated genes related to synapse and neurodevelopment included *Pax6*, which directly binds and activates the promoter of somatostatin^119,120^, the somatostatin receptor *Sstr3*, the axon guidance genes *Robo2*, as wells as the IEGs *Jun*, *Junb*, *Jund*, *Fos*, *Fosb* and *Npas4* (Figure 9H; green). Notable genes related to ER stress and protein folding included downregulation of Hsp70 family members *Hspa8*, *Hspa1b*, and *Hspa1a*, as well as co-chaperones of Hsp40 *Dnaja1*, *Dnaja4*, and *Dnajb2* (Figure 9H; grey).

## Discussion

Individuals with Down syndrome have an increased susceptibility to developing T2D. However, the etiology of T2D in Down syndrome is not fully understood^3^. While environmental factors, like higher rates of obesity^5^ and reduced physical activity^121^ in people with Down syndrome compared to the general population could explain some of this phenomenon, it is increasingly appreciated that direct genomic alterations due to trisomy 21 contribute to the increased risk of T2D in Down synrome^3^. Recent studies have identified underlying defects in systemic metabolism in people with Down syndrome as well as in mouse models of trisomy 21, linked to genome-wide gene expression dysregulation in the liver, muscle, hypothalamus, and adipose tissues^6–8^. However, the role of genomic dysregulation in the islets of Langerhans as a causative driver of T2D in Down syndrome has not been explored. Here, we leverage a mouse model of Down syndrome, Ts65D, to explore the hypothesis that innate transcriptional alterations in the islets due to trisomy 21 increase the likelihood of developing T2D in individuals with Down syndrome. Our results provide a comprehensive transcriptomic atlas of the islet of Langerhans, describing organ- and cell-type level gene expression differences caused by trisomy at the syntenic region of human chromosome 21, highlighting transcriptional links with T2D susceptibility in individuals with Down syndrome.

We corroborate previous reports^8,15^ that young adult (6-8 weeks old) trisomic Ts65Dn mice are glucose intolerant, despite having normal insulin tolerance, implying islet dysfunction. Interestingly, islets of trisomic Ts65Dn mice have mild alterations in their endocrine cell types proportions (with less β cells and more α cells). However, the overall β cell area was similar between islets of trisomic and disomic mice, suggesting some degree of compensatory β cell hypertrophy in the trisomic islets. Importantly, these results are consistent with a previous report by Butler *et al*.^122^, showing no difference in the average insulin-immunoreactive area between a cohort of non-diabetic humans with Down syndrome and age-matched controls.

To gain insight into islet defects in Down syndrome, we performed scRNA seq on isolated islets from trisomic Ts65Dn mice and disomic controls. DEGs in trisomic islets are not restricted to the triplicated region but extend across the entire genome, in agreement with human transcriptomic studies^123–126^. Moreover, we saw both upregulation and downregulation of large numbers of genes, indicating a global gene expression dysregulation in trisomic islets. We show that this genome-wide dysregulation generates different transcriptional signatures depending on cell type and sex. However, despite these differences, several common transcriptional themes were observed between most cell types and both sexes, that can contribute to the unique etiology of Down syndrome-related T2D.

The genomic dysregulation in Down syndrome takes effect at multiple stages across the lifespan, from interfering with embryonic and postnatal development, to negatively affecting cellular adaptation to environmental stresses^127^. We observed the expected ∼1.5-fold over-expression of several genes located on the triplicated segment of trisomic Ts65Dn mice, and whose misexpression was previously reported to affect endocrine cells differentiation, function, or adaptation to diabetogenic stresses. For example, we detected overexpression of *Dyrk1a* and *Rcan1*, two triplicated genes overexpressed in most endocrine cells, which is expected to reduce the proliferation capacity in response to higher demand for insulin^24–27^, via inhibition of the calcineurin-NFAT pathway^128–130^; overexpression of *Rcan1* has also been implicated in mitochondrial dysfunction in β cells in Down syndrome^15,27^. Higher levels of *Mpc1* and *Sod1* expression, two other triplicated genes overexpressed in most endocrine cells examined in our study, are similarly expected to negatively affect mitochondrial health and oxidative stress, respectively^32,33^. Overexpression of the triplicated gene, *Bach1*, a transcriptional repressor of antioxidative enzymes^31^, is expected to aggravate this effect. Indeed, cells from individuals with Down syndrome and from Down syndrome mouse models were reported to show high levels of mitochondrial dysfunction and elevated oxidative stress^8,11,15,20,27,131,132^. This phenomenon is particularly important to β cells, which are exceptionally vulnerable to mitochondrial dysfunction and oxidative stress due to the co-occurrence of high production of reactive oxygen species, low expression of antioxidizing enzymes, and reliance on mitochondrial oxidative metabolism for insulin secretion^133–135^. Finally, the overexpression of the amyloid precursor, *App*, and the protease *Bace2*, in our dataset, is expected to increase the deposition of amyloid plaques^28–30^, consistent with a previous report describing increased amyloid plaques in islets of individuals with Down syndrome^11^.

Beyond the direct effects of triplicated genes, our study revealed altered expression of hundreds of non-triplicated genes that may contribute to the increased susceptibility of individuals with Down syndrome to developing T2D, through both cell-autonomous effects and disruption in intra-islet communication among endocrine cell types. One such alteration, shared among all endocrine cell types and between males and females, is the robust down-regulation of many IEGs, such as *Jun*, *Fos*, *Myc*, *Egr1*, and *Npas4*. IEGs play important roles in the islet by acting as rapid responders to various stimuli like glucose, metabolic and inflammatory stresses, incretin hormones, and more, and by influencing adaptive mechanisms like hormone secretion, cell survival, and cell proliferation^53–55,69,136,137^. The reduction of IEGs in islets of ALT1 mice suggests a reduced adaptive response to metabolic changes, reduced secretory activity, reduced incretin signaling in the islet, or a combination of these. Another notable example is the misexpression of many ER stress and protein folding proteins. Chaperones from the Hsp40 family (*Dnaja1*, *Dnaja4*, *Dnajb2*, *etc*.) and from the Hsp70 family (*Hspa8*, *Hspa1b*, *Hspa1a*, *etc*.), which were notably downregulated in all endocrine cells of both male and female ALT1 mice; however, other ER stress proteins, like the misfolded proteins ERAD genes *Selenos*, *Edem2*, *Manf*, and others, were upregulated in β and δ cells, suggesting that the cells are unable to activate a coordinated response to ER stress.

Our single cell transcriptomic profile also highlights many alterations that could adversely affect cellular communications at the whole-islet level, the most notable of which is arguably the reduced expression of critical genes involved in incretin signaling. *Gipr*, the receptor for the incretin hormone GIP, was significantly downregulated in β cells of both male and female ALT1 mice; *Btg2*, which mediates GLP-1 signaling in the islet, was also significantly downregulated in α cells and β cells of both sexes. Importantly, brains of trisomic Ts65Dn mice show downregulation of *Glp1r*^138^, suggesting that defects in incretin signaling may have a systemic effect on Down syndrome-related T2D. Importantly, we found a downregulation of Notch pathway genes (*Notch1*, *Dll4*) in β cells of both male and female ALT1 mice indicating aberrant intra-islet communication in Down syndrome. Notch signaling play important roles in endocrine cell development, β cell function, and β cell survival under diabetogenic stresses^139–145^. Additionally, sex-dependent dysregulation of somatostatin receptors *Sstr2* and *Sstr3* (upregulated in male ALT1 α cells, downregulated in female ALT1 α cells) could account for some of the sexually dimorphic phenotypes seen in ALT1 mice.

In summary, our work provides the first comprehensive single-cell transcriptomic atlas of the islets of Langerhans in a mouse model of Down syndrome. Our dataset revealed cell-type and sex-specific transcriptional signatures and identified a transcriptional program associated with increased susceptibility to T2D. Our data show that trisomy negatively affects the islets at multiple levels, either directly by overexpressing triplicated genes, or indirectly via genome-wide gene expression dysregulation. Our work thus puts forth the islets of Langerhans as a potential driver of Down syndrome-related T2D and provides a comprehensive resource for the field to further dissect specific mechanisms and potential therapeutic targets for this important yet understudied comorbidity of Down syndrome.

## Limitations of the study

While mouse models such as Ts65Dn and Dp(16)1Yey are valuable for the study of Down syndrome-related T2D by assessing whole-body metabolism and inter-tissue phenotypes, they are limited in the information they can provide on the direct, islet-specific consequences of trisomy 21 in two critical aspects: 1) due to the genomic alteration in these mice being present in all cells of the animal, it is difficult to discern which islet phenotypes are a direct result of the genomic alteration, and which phenotype are indirectly caused by the islets’ response to secondary metabolic changes in other tissues; 2) in mice, the genomic regions syntenic to human chromosome 21 are spread over three different mouse chromosomes: chromosome 16, chromosome 17, and chromosome 10. This confounds conclusions about phenotypes related to structural genomic abnormalities in human Down syndrome cells. As pancreas and islet samples from individuals with Down syndrome are extremely scarce, further research would benefit from using human trisomy 21 iPSC lines^20,146^, which, combined with state of the art protocols for differentiating β cells-, α cells-, and δ cells-enriched stem-cells-derived islet-like clusters^147–150^, will be essential in modeling human-specific genomic dysregulation in Down syndrome in the islets of Langerhans, and how it affects Down syndrome-related T2D.

## Materials and Methods

### Animals

All procedures were approved by the University of Wisconsin-Madison Institutional Animal Care and Use Committee (Protocol #M005221) and conducted in accordance with the NIH Guide for the Care and Use of Laboratory Animals. Trisomic Ts65Dn mice (B6EiC3Sn.BLiA-Ts(17^16^)65Dn/DnJ; The Jackson Laboratory, Stock #005252; RRID:IMSR_JAX:005252) and euploid littermate controls were obtained from The Jackson Laboratory Cytogenetic and Down Syndrome Models Resource. The colony is maintained at The Jackson Laboratory by crossing trisomic Ts65Dn females to B6EiC3Sn.BLiAF1/J males (JAX Stock #003647). All animals were genotyped at The Jackson Laboratory prior to shipment. Genotype was reconfirmed upon arrival in our facility by PCR amplification of the Ts(17^16^)65Dn translocation breakpoint junction using the Ts(17<16>;EIF1AX-Xist)67Jlaw separated PCR assay (JAX Protocol 24762).

### Oral Glucose Tolerance Tests

Oral glucose tolerance tests were performed on 6-week-old ALT1 and WT mice. Mice were fasted for 6 hours beginning in the morning and then administered glucose at a dose of 2 g/kg body weight via oral gavage. Blood glucose was measured from the tail tip at 0, 15, 30, 45, 60, 90, and 120 minutes post-gavage using a Contour Next EZ Blood Glucose Meter (Ascensia Diabetes Care).

### Insulin Tolerance Test

Insulin tolerance tests were performed on independent cohorts of 8-week-old mice. Mice were fasted for 4 hours beginning in the morning and then administered Humulin R (Lilly, U-100) intraperitoneally at a dose of 1 U/kg for males and 0.5 U/kg for females. Blood glucose was measured from the tail tip at 0, 15, 30, 45, 60, 90, and 120 minutes post-injection using a Contour Next EZ Blood Glucose Meter.

### Islet Histology and Immunofluorescence

Mice were euthanized with CO_2_, and their pancreata were dissected out and fixed with 4% paraformaldehyde (PFA) for 3 hours on a rotator at 4°C. Following fixation, tissues were incubated in a 30% (w/v) sucrose solution in PBS and then embedded in OCT compound (FSC 22 Blue, Cat #3801481). Cryosections were cut at 10 µm thickness on a Leica CM 1860 cryostat. Two sets of 5 slides were collected per pancreas, with 200 µm between sections. Sections were processed using the Mouse on Mouse (M.O.M.™) Basic Kit (Vector Laboratories, Cat #BMK-2202) according to the manufacturer’s protocol with slight modifications, as described in^151^. Primary antibodies were applied overnight at 4°C: guinea pig anti-insulin (Dako, Cat #IR002, 1:6), mouse anti-glucagon (Sigma Aldrich, Cat #G2654-100UL, 1:200), and rabbit anti-somatostatin (Phoenix Pharmaceuticals, Cat #G-060-03, 1:500). Secondary antibodies were applied for 1 hour at room temperature: Alexa Fluor 488 donkey anti-rabbit (Jackson Immunoresearch, Cat #711-546-152, 1:500), Alexa Fluor 594 donkey anti-guinea pig (Jackson Immunoresearch, Cat #706-585-148, 1:500), and Alexa Fluor 647 donkey anti-mouse (Jackson Immunoresearch, Cat #715-605-150, 1:500). Nuclei were counterstained with DAPI (Sigma Aldrich, Cat #D9542-5MG, 1:1000 from a 0.5 mg/mL stock) overnight.

### Imaging and Quantification

Confocal images were acquired on a Nikon AXR Confocal Microscope at the Optical Imaging Core, University of Wisconsin–Madison (Grant #1S10O34394-01), using a 40×/1.3 NA oil immersion objective and NIS Elements Viewer software. Ten islets were quantified per mouse (n = 4 mice per genotype; 9 weeks old). Cell type proportions were analyzed using QuPath v0.6.0 by annotating cells based on channel intensities corresponding to glucagon, insulin, and somatostatin, respectively. Data were obtained by dividing the number of positive cells for each cell type by the total number of nuclei detected. Beta cell area was analyzed in ImageJ/FIJI v1.54f by outlining the insulin-marked area using the Color Threshold function on the Analyze panel, with the RenyiEntropy thresholding method. A minimum of 40 DAPI-positive cells was required to define an islet unit. Groups were compared using unpaired t-tests in GraphPad Prism; p < 0.05 was considered significant. Analyses were not blinded.

### Islet Isolation

Mice were perfused through the common bile duct with a collagenase solution containing 0.5 mg/mL Collagenase P (Roche, Cat #11249002001) in RPMI 1640 (Corning, Cat #10-040-CM) supplemented with 1× Penicillin/Streptomycin (P/S; Corning, Cat #30-002-CI). Islets were purified by density gradient centrifugation using Histopaque 1077 and 1119 (Sigma Aldrich, Cat #10771 and 11191, respectively), followed by manual hand-picking under a stereomicroscope. Isolated islets were maintained in RPMI 1640 supplemented with 10% FBS (Cytiva, Cat #SH30088.02) and P/S at 37°C and 5% CO2. For scRNA seq experiments, islets were allowed to equilibrate for 1 hour at 37°C and 5% CO2 before dissociation. For GSIS experiments, islets were allowed to recover overnight.

### Single-Cell RNA Sequencing

Following equilibration, isolated islets were dissociated into single-cell suspensions using TrypLE Express (Gibco, Cat #12604021) for 10 minutes at 37°C. Cell viability and concentration were assessed using a LUNA-FX7 Automated Cell Counter (Logos Biosystems) prior to loading onto the 10x Genomics Chromium controller. Following library construction, library quality was verified by shallow sequencing on an Illumina MiSeq prior to full-depth sequencing on the NovaSeq X Plus. Single-cell libraries were prepared using the 10x Genomics Chromium platform with the Single Cell 3’ GEM-X v4 chemistry, targeting approximately 10,000 cells per sample. Libraries were sequenced on an Illumina NovaSeq X Plus 10B with paired-end 2 × 50 bp reads, targeting approximately 50,000 reads per cell. Sequencing was performed at the University of Wisconsin Biotechnology Center Gene Expression Center. Raw sequencing reads were demultiplexed and aligned to the GRCm39 (mm39) mouse reference genome (build 2024-A) using Cell Ranger v8.0.1 (10x Genomics). Cell Ranger generated filtered feature-barcode matrices for each sample, yielding a total of 84,410 cells across 9 samples prior to downstream quality control filtering (range: 4,322–12,719 cells per sample; mean reads per cell: 37,363–57,218).

### RNA seq Quality Control and Filtering

All downstream analyses were performed in R v4.4.2 using Seurat v5.4.0. Ambient RNA contamination was first estimated and corrected for each sample independently using SoupX v1.6.2, with automated contamination fraction estimation (*autoEstCont*) and count adjustment (*adjustCounts*). The effectiveness of ambient RNA removal was validated by inspecting marker-specific correction maps for Ins1 and Mafa. Potential doublets were then identified using scDblFinder v1.20.2 with cluster-based artificial doublet simulation (*clusters = TRUE*). Quality control filtering was applied with the following thresholds: cells were retained if they expressed between 2,500 and 8,000 genes (*nFeature_RNA*), had less than 5% mitochondrial transcript content (identified by the mt-prefix), and received a doublet score below 0.5. After quality control filtering, 45,321 cells were retained across the nine samples (ALT1-FM-001: 5,950; ALT1-FM-002: 5,305; ALT1-M-001: 6,176; ALT1-M-003: 5,600; WT-FM-002: 3,810; WT-FM-004: 2,913; WT-M-001: 4,277; WT-M-002: 5,214; WT-M-004: 6,076).

### RNA seq Normalization, Integration, and Clustering

Filtered count matrices were normalized using log-normalization with a scale factor of 10,000 (Seurat *NormalizeData*). The top 2,000 highly variable features were identified using variance stabilizing transformation (VST; Seurat *FindVariableFeatures*), and expression values were scaled across all genes (*ScaleData*). Principal component analysis (PCA) was performed, and sample-level batch effects were corrected using Harmony v1.2.4, applied to the PCA embedding via *IntegrateLayers* (method = HarmonyIntegration). A shared nearest-neighbor (SNN) graph was constructed from the first 10 Harmony-corrected dimensions, and cells were clustered using the Louvain algorithm. Multiple clustering resolutions were evaluated (0.25, 0.3, 0.35, 0.37), and a final resolution of 0.37 was selected based on concordance between cluster boundaries and known marker gene expression patterns, yielding 16 transcriptionally distinct clusters (numbered 0–15). Uniform Manifold Approximation and Projection (UMAP) was computed on the Harmony-corrected dimensions (dims 1:10, seed = 7) for visualization.

### Cell Type Annotations

Cell type identities were assigned to each cluster based on the expression of canonical marker genes. Cluster-level marker genes were identified using Seurat FindAllMarkers (only positively enriched genes; avg_log2FC > 1). The following marker genes were used for annotation: β cells (*Ins1*, *Ins2*, *MafA*, *Ucn3*); α cells (*Gcg*, *Irx2*, *Arx*); δ cells (*Sst*, *Hhex*, *Dscam*); PP cells (*Irx2*, *Ppy*); ductal cells (*Krt19*, *Hnf1b*); endothelial cells (*Pecam1*, *Cdh5*); immune cells (*Ptprc*, *Cd68*); pericytes (*Pdgfrb*, *Aspn*); mesenchymal (*Sostdc1*, *Ptn*, *Sfrp1*); and fibroblasts (*Pdgfra*, *Vim*, *Col3a1*). In total, 13 cell type identities were assigned: α, β, δ, two polyhormonal populations (Poly_1, Poly_2), PP, Ductal, Endothelial, Endo-Peri, Pericytes, Fibroblasts, Immune, and Mesenchymal. Hierarchical clustering (Ward.D2 linkage on Euclidean distances of cluster-averaged expression profiles) was used to assess transcriptomic relationships between clusters.

### Differential Gene Expression Analysis

Differential gene expression (DE) analysis was performed at the single-cell level using the Wilcoxon rank-sum test with Bonferroni correction as implemented in Seurat *FindMarkers* (with accelerated computation via presto v1.0.0). For each cell type, trisomic (ALT1) cells were compared to disomic (WT) controls, with analyses performed separately by sex (male and female). Genes were considered differentially expressed if they met a threshold of |log_2_FC| ≥ 0.25 and a Bonferroni-adjusted p-value < 0.05.

### Gene Ontology Enrichment Analysis

Gene Ontology (GO) enrichment analysis was performed using clusterProfiler v4.14.6 with the *enrichGO* function and the org.Mm.eg.db v3.20.0 annotation database. Enrichment was tested against a background of all expressed genes detected in the dataset. P-values were adjusted for multiple testing using the Benjamini-Hochberg method, with a false discovery rate threshold of q < 0.05 (clusterProfiler default settings). While all three GO categories were analyzed (Biological Process, Molecular Function, and Cellular Component), results presented in this study focus on Biological Process (BP) terms. Redundant GO terms were reduced using the built-in *simplify* function in clusterProfiler (⍺ = 0.7), and the top 20 enriched BP terms were visualized as dot plots. To organize enriched terms into biologically coherent groups for discussion, terms were first clustered by Jaccard similarity, followed by iterative centroid-based refinement, and then manually curated to reassign terms that were misclassified or not captured by the automated clustering. This organizational step was performed solely to facilitate the presentation and interpretation of results and did not alter the underlying enrichment statistics or significance calls.

### Chromosomal Distribution of Differentially Expressed Genes

Differentially expressed genes were mapped to their chromosomal positions using gene annotation from biomaRt v2.62.1 and org.Mm.eg.db. The distribution of DE genes across mouse chromosomes 1–19, X, and Y was summarized per cell type, separated by sex and direction of change (upregulated vs. downregulated). Genes mapping to the triplicated segment were specifically flagged.

### Statistical Analysis

oGTT and ITT time-course data were analyzed using two-way ANOVA with Šídák’s multiple comparisons test. oGTT area under the curve (AUC) was analyzed using an unpaired t-test. Cell proportion comparisons (histology) were performed using unpaired t-tests. For all metabolic and histological analyses, a p-value < 0.05 was considered statistically significant. Statistical tests for these analyses were performed using GraphPad Prism. For scRNA seq differential expression, the Wilcoxon rank-sum test with Bonferroni correction was used as described above.

### Software and Data Availability

All single-cell RNA sequencing analyses were performed in R v4.4.2 using Seurat v5.4.0 (SeuratObject v5.3.0). Key additional packages included: Harmony v1.2.4 (batch integration), SoupX v1.6.2 (ambient RNA correction), scDblFinder v1.20.2 (doublet detection), clusterProfiler v4.14.6 (GO enrichment), org.Mm.eg.db v3.20.0 (gene annotation), presto v1.0.0 (accelerated DE testing; GitHub: immunogenomics/presto), biomaRt v2.62.1 (chromosomal mapping), MAST v1.32.0, DESeq2 v1.46.0, and ggplot2 v4.0.1 with patchwork v1.3.2 and viridis v0.6.5 for visualization. GO dot plots, volcano plots, UMAP projections, and chromosomal distribution maps were generated using custom visualization scripts adapted from scRNAVis (Ref.^152^). Additional custom R scripts were used for Jaccard similarity-based GO term clustering, and chromosomal distribution analysis. Confocal image analysis was performed using ImageJ/FIJI v1.54f and QuPath v0.6.0. Statistical analyses for metabolic and histological data were performed using GraphPad Prism. RNA seq data and code were deposited in GEO and GitHub under the accession number XXXXXX and URL XXXXXXX, respectively.

## Acknowledgements

We thank past and present members of the Blum and the Lo Sardo labs for valuable discussions. We thank Dr. Anita Bhattacharyya for critical reading and comments on the manuscript. We thank Youngha Kim and the University of Wisconsin Biotechnology Center Gene Expression Center for library preparation and sequencing services. We also thank Dr. Anthony Veltri of the UWBC Bioinformatics Resource Center for his expert assistance with read alignment and data preprocessing. The sequencing and bioinformatics experiments were supported in part by the UW-Madison Office of the Vice Chancellor for Research and Graduate Education and utilized instrumentation supported by the NIH Shared Instrumentation Grant S10 OD025052. This work was funded in part by grants from the NIH (R01DK131438, R01DK121706) and the Jerome Lejeune Foundation (GRT-2022A/2123) to BB, The UW-Madison ICTR Pilot Award to BB and VLS.

---

**Supplementary Figure 1.**
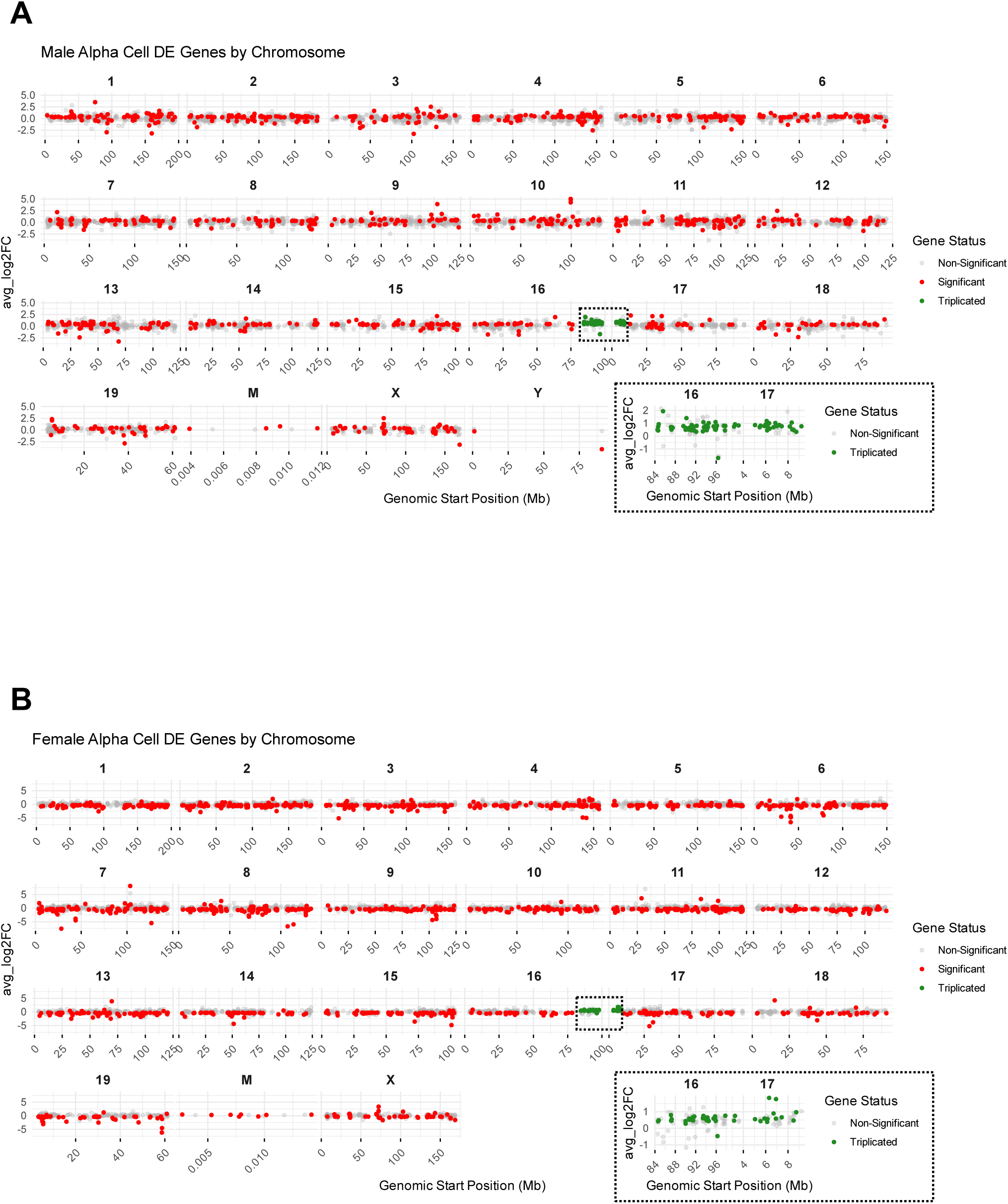
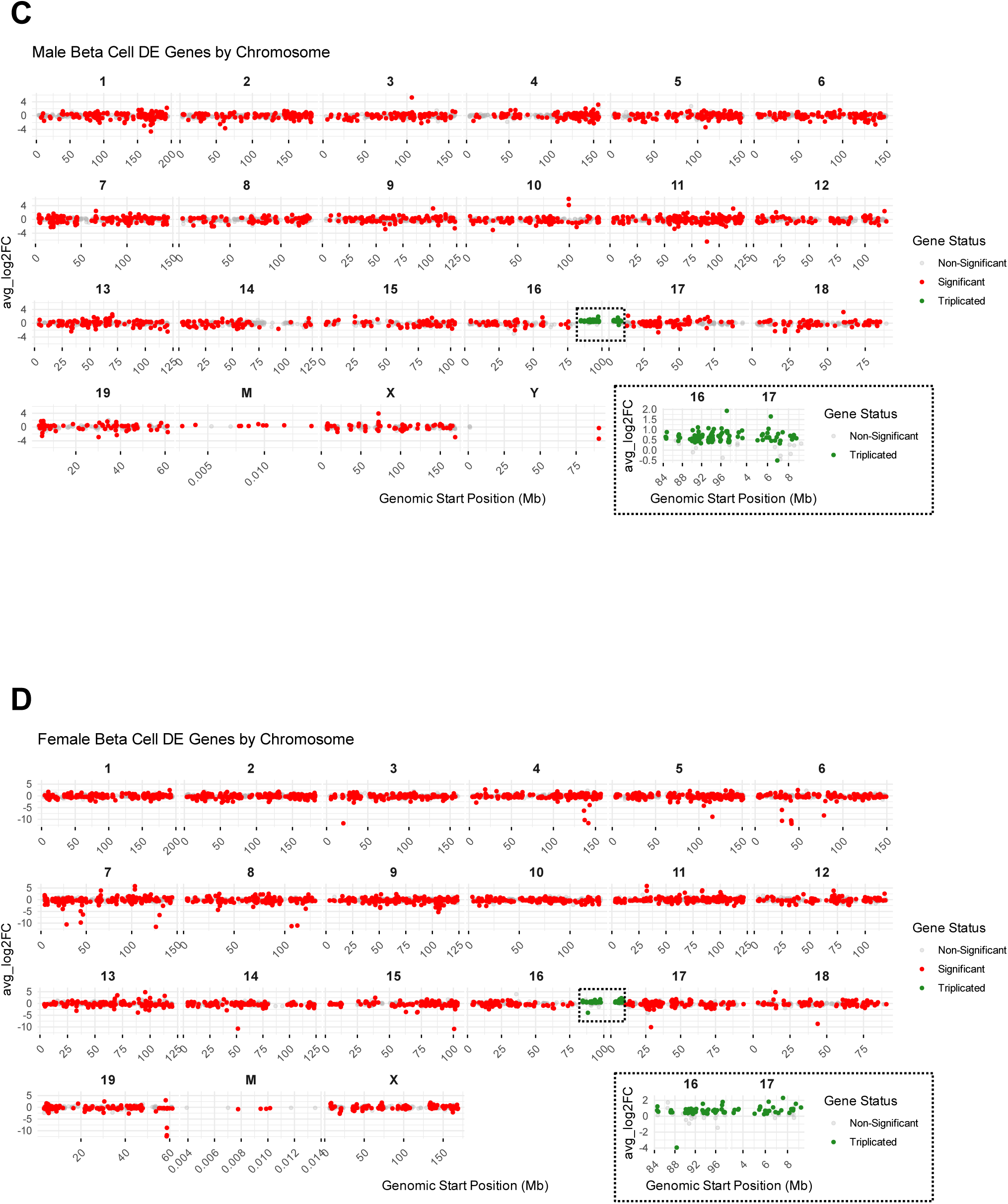
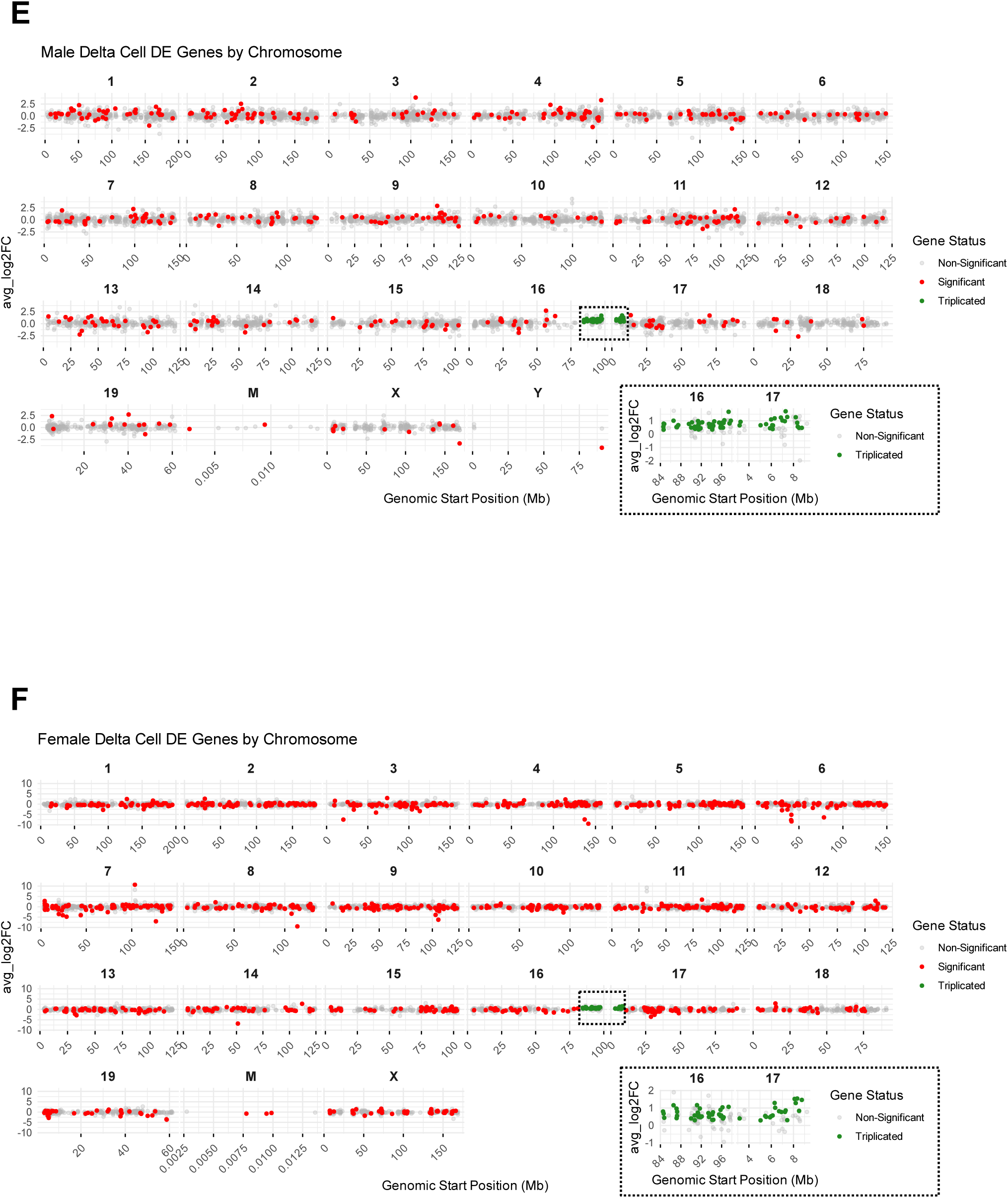
Chromosomal distribution of misexpressed genes by endocrine cell type. **(A-F)** Plots of differentially expressed genes by chromosomal position in male α cells **(A)**, female α cells **(B)**, male β cells **(C)**, female β cells **(D)**, male δ cells **(E)**, and female δ cells **(F)**. The x-axis denotes genomic location, and the y-axis denotes average log₂ fold-change. Red = significantly misexpressed (upregulated or downregulated), green = misexpressed genes within the triplicated region, grey = non-significant genes.

